# Dynamic ventral disc contraction is necessary for *Giardia* attachment and host pathology

**DOI:** 10.1101/2023.07.04.547600

**Authors:** C Nosala, KD Hagen, SL Guest, NA Hilton, A Müller, M Laue, C. Klotz, A Aebischer, SC Dawson

## Abstract

*Giardia lamblia* is a common parasitic protist that infects the small intestine and causes giardiasis, resulting in diarrhea, vomiting, weight loss, and malabsorption. Giardiasis leads to cellular damage, including loss of microvilli, disruption of tight junctions, impaired barrier function, enzyme inhibition, malabsorption, and apoptosis. In the host, motile *Giardia* trophozoites attach to the duodenal microvilli using a unique microtubule organelle called the ventral disc. Despite early observations of disc-shaped depressions in microvilli after parasite detachment, little is known about disc-mediated attachment mechanisms and there little direct evidence showing that parasite attachment causes cellular damage. However, advancements in *in vitro* organoid models of infection and genetic tools have opened new possibilities for studying molecular mechanisms of attachment and the impact of attachment on the host. Through high-resolution live imaging and a novel disc mutant, we provide direct evidence for disc contraction during attachment, resolving the long-standing controversy of its existence. Specifically, we identify three types of disc movements that characterize contraction, which in combination result in a decrease in disc diameter and volume. Additionally, we investigate the consequences of attachment and disc contractility using an attachment mutant that has abnormal disc architecture. In a human organoid model, we demonstrate that this mutant has a limited ability to break down the epithelial barrier as compared to wild type. Based on this direct evidence, we propose a model of attachment that incorporates disc contraction to generates the forces required for the observed “grasping” of trophozoites on the host epithelium. Overall, this work highlights the importance of disc contractility in establishing and maintaining parasite attachment, leading to intestinal barrier breakdown.

## Introduction

*Giardia lamblia* is a common zoonotic protistan parasite that colonizes the small intestine, causing the diarrheal disease giardiasis. Giardiasis occurs worldwide and is especially prevalent where water quality is poor [1]. Early and recurrent childhood *Giardia* infection can exacerbate malnutrition and cause development delays [2]. Hosts ingest *Giardia* cysts, which then excyst to release flagellated trophozoites that colonize and attach to the duodenal microvilli with the “ventral disc”, a conspicuous cup-like microtubule (MT) organelle unique to *Giardia* species [3]. Common symptoms of *Giardia* infection include diarrhea, vomiting, weight loss, and malabsorption [2]. At the cellular level, the consequences of *Giardia* infection are reported to include damage to and loss of enterocyte microvilli, disrupted tight junctions and barrier function, inhibition of brush border enzymes, malabsorption, and apoptosis [2].

The pathophysiological consequences of *Giardia* infection have traditionally been attributed to indirect, rather than direct interactions of the trophozoite with the host epithelium during parasite attachment [2]. These indirect interactions include factors such as the unique substrates and products of parasite metabolism, secretion of cysteine proteases, or disruptions to the host microbiome [2,4]. The intricate interplay between parasite attachment, colonization, and direct damage to the epithelium was a key component of early observations of *Giardia*-host interactions, and researchers specifically noted the presence of ventral disc-shaped depressions that remained in the microvilli after parasites detached from the epithelium [5,6] [6].

Initially, actin-based mechanisms of the disc-associated lateral crest, rather than movements of the disc itself, were proposed to generate the underlying biophysical forces required for the grasping behavior of trophozoites on the epithelium [7], and lateral crest mediated attachment of the disc was assumed to either directly cause tissue damage or indirectly limit absorption of nutrient in the small intestine. Thus actin-based disc contractility has since been generally discounted. In contrast, the oft-cited hydrodynamic suction model of Holberton [8] introduced an innovative hypothesis for attachment that discounted contractile forces of the disc [6] and instead proposed that beating of the ventral flagella produced a hydrodynamic current drawn underneath a rigid, inflexible disc. Using flagellar motility mutants, we challenged this hypothesis [9], and demonstrated that attachment to surfaces was unaffected when flagellar motility was impaired. Thus, ventral flagellar beating alone is not necessary for maintaining hydrodynamic suction.

In the absence of either actin-based contraction of the lateral crest [7] or hydrodynamic suction mediated by flagellar motility, the disc itself becomes the obvious candidate for force generation for attachment. Consequently, a thorough comprehension of disc architecture is essential to unravel attachment mechanisms and understand their impacts on the host. Attachment mechanisms likely derive from the novel structural components or protein complexes comprising the disc. As an organelle of attachment, the ventral disc is an elaborate domed structure defined by approximately 100 parallel MTs that spiral until they overlap in the “overlap zone”, a region where the upper and lower disc layers are connected (Figure 1A and [3]). The spiral MTs scaffold additional unique protein complexes including the ribbon-like microribbon-crossbridge (MR-CB) complex. The asymmetric trilaminar microribbons track the entire MT spiral and extend 400 nm into the cytoplasm, while the crossbridges connect parallel microribbons. Additional protein complexes comprise the disc perimeter, or “disc margin,” and the lateral crest, which is an additional repetitive structure that lies outside the MT spiral on the disc circumference and been predicted to grasp the host microvilli [6]. We previously identified close to 90 disc-associated proteins (DAPs) that associate with specific disc substructures, such as the lateral crest or overlap zone [10], that could be involved in force generation for attachment (Figure 1). With respect to the role of DAPs in disc structure, we have recently used both CRISPR-based knockdowns and CRISPR-based quadruple KOs to show that some DAPs are critical for the disc domed architecture[11,12], and others are essential in providing for disc MT stability and overall disc hyperstability [10].

**Figure 1.**
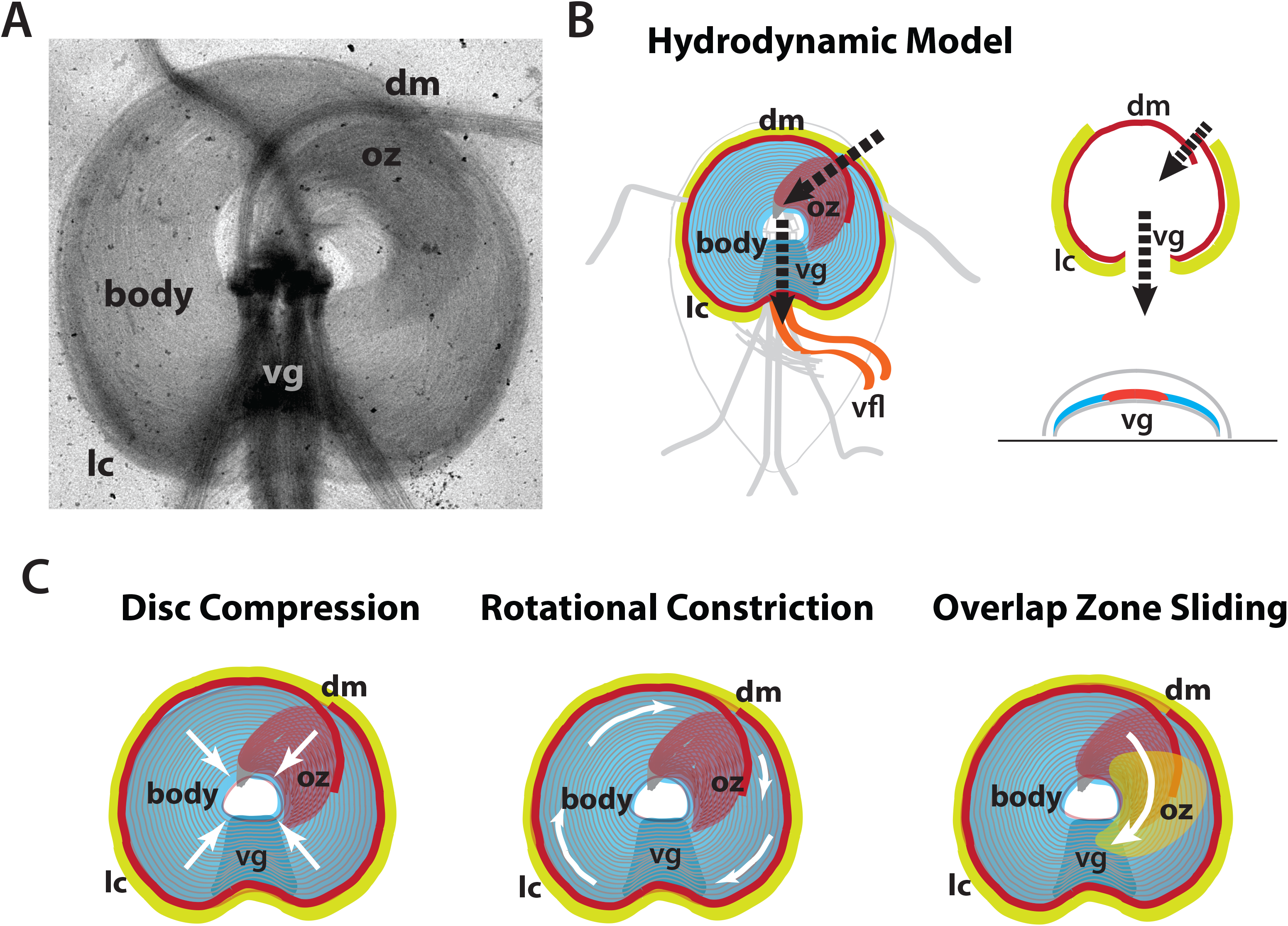
Three types of disc movements may be associated with disc contraction. As illustrated in the negative stained disc preparation (A) the ventral disc microtubules (body) are nucleated from the basal bodies. Distinct structural regions of the disc and lateral crest (lc) are noted, including the ventral groove (vg), the disc margin (dm), and the overlap zone (oz). In B, hydrodynamic currents generated by the ventral flagellar beating (vfl) and fluid flowing through a gap in the disc margin (dm) underneath the ventral groove (vg) were proposed to generate suction-based attachment. In C, three different types of movements associated with specific disc structural elements have also been predicted to generate contractile forces that mediate attachment. Microribbon-crossbridge complex compression in “disc compression” movements; the rotational constriction of the lateral crest; or the rotation and sliding of the upper and lower layers of the disc in the overlap zone region.

Combined with new *in vitro* models such as human gastrointestinal epithelium iPSC organoids that recapitulate epithelial damage and barrier function, the advent of new genetic tools to create stable *Giardia* attachment mutants offers promise for discovering the impact of trophozoite attachment and colonization on the host. We recently developed robust CRISPR-based genetic tools to create highly penetrant disc attachment null mutants, and have used these nulls with the organoid infection model to test the impact of attachment [11,13]. Our results indicated that disc-mediated attachment is necessary for the disruption of epithelial barrier integrity and functioning[11]. Nonetheless, with respect to cellular mechanisms of attachment, we still lack direct evidence for the disc contractility initially reported [6] and we also lack direct evidence that disc-mediated attachment and/or contractile functions cause cellular damage to the host epithelium [2].

Yet to paraphrase the classic maxim: the ***absence of evidence*** for disc contraction ***is not evidence of the absence*** of disc contraction. Here, using high resolution live imaging of disc dynamics with DAPs that mark the lateral crest, we provide direct evidence for disc contractility, resolving the long-standing controversy regarding disc contraction during parasite attachment. Live imaging of disc “marker” strains allowed us to track and quantify movements of disc components relative to each other in seconds in live attaching trophozoites. We define three types of disc and lateral crest movements that characterize overall disc contraction that decreases the disc diameter by about 0.5 µm and disc volume by roughly 10-15%. Further, we test the consequences of attachment and disc contractility for the first time, using a DAP quadruple null mutant (DAP7268) that lacks contractility and has an aberrant disc “Baumkuchen” architecture with multiple disc overlap zones and lateral crests. Using the infections with human organoid derived monolayers [13] we show that Baumkuchen mutants have a limited ability to cause epithelial barrier breakdown. Based on this new, direct empirical molecular genetic evidence, we propose a model of attachment with a mechanism for disc contraction that could generate forces for the grasping behavior initially observed over 50 years ago. Overall, this work demonstrates that disc contractility is necessary for establishing and maintaining parasite attachment and that such contractility-based attachment is necessary for intestinal barrier breakdown.

## Materials and Methods

### Giardia cultivation, electroporation, and isolate cloning

All strains used in this study, including tagged, knockout, and other variants, were derived from *Giardia lamblia* WBC6 strain ATCC 50803. Strains were thawed from frozen stocks and cultivated at least 24 to 48 hours prior to phenotypic analysis unless otherwise specified. Routine cultivation was carried out in 16-ml screw-cap plastic tubes (BD Falcon) without agitation. TYI-S-33 growth medium was supplemented with bovine bile, 5% adult bovine serum, and 5% fetal bovine serum, and the cultures were maintained at 37°C. Upon reaching confluency, culture tubes were incubated on ice for 30 minutes, and 0.5 to 1 ml of detached trophozoites was added to 12 ml of pre-warmed medium. Antibiotics were added into the culture medium when selecting for episomal plasmids (GFP-tagged strains) or gene knockouts ([11] and further details provided below). Electroporation of DNA constructs (plasmids or linear homology-directed repair (HDR) templates) into *Giardia* were performed under conditions described previously [12], using 20 to 40 μg DNA and 1 x 10^7^ trophozoites per electroporation. Electroporated cells were then incubated in standard culture tubes at 37C, with medium replacement every 48 hours. Lower antibiotic concentrations were used for initial selection and antibiotics were increased to maintenance levels once culture confluency reached >50%. Concentrations used were: puromycin (10 µg/ml and 50 µg/ml), blasticidin (75 µg/ml and 150 µg/ml), and hygromycin B Gold (600 µg/ml and 1200 µg/ml). All antibiotics were purchased from Invivogen except G418 (Fisher Scientific).

Clonal cell lines were obtained by limiting dilution. Iced, detached trophozoites were counted with a Luna-II cell counter (Logos Biosystems). Trophozoites were diluted to 2 cells per ml in fresh TYI-S-33 containing antibiotics used for selection, and 250 μl aliquots (0.5 cells/well) were added to 96 well tissue culture plates (BD Falcon). Plates were incubated at 37C in Mitsubishi AnaeroPack 2.5L rectangular jars with a Mitsubishi AnaeroPack-Anaero Gas Generator sachet (ThermoScientific) for 7 to 10 days. Typically, 20 to 30% of plate wells contained cells, and most wells with cells had reached 80 to 100% confluency. Plates were iced 30 minutes and detached trophozoites were transferred to 8 ml screw-capped tubes (BD Falcon). Once these tubes reached >50% confluency, cells were propagated in standard 12 ml tubes.

### Design and creation of DAP7268 and DAP7268_12139mNG quadruple knockout (null) strains

Our CRISPR quadruple knockout strategy has been described elsewhere [11]. Briefly, a guide RNA targeting the coding region of DAP7268 at position 908 (gRNA908R, 5’-AGTTGATCCGGAACGCTGTG-3’) was selected using Benchling’s CRISPR ‘Design and Analyze Guides’ tool (https://benchling.com/crispr) with an NGG PAM sequence and the *Giardia lamblia* ATCC 50803 genome (GenBank Assembly GCA_000002435.1). A slightly modified vector, Cas9U6g1pac, was used to express both the gRNA and Cas9 fused to the *Giardia* GL50803_2340 NLS—the gRNA expression cassette in this new vector differs from our previously published CRISPR and CRISPRi vectors [11,14] in that 12 bases encoding U6 snRNA that were immediately downstream from the U6 promoter have been removed. Complementary gRNA oligos (5’-cggcAGTTGATCCGGAACGCTGTG-3’ and 5’-aaacCACAGCGTTCCGGATCAACT-3’) with 4 base overhangs were annealed and ligated to BbsI-digested Cas9U6g1pac as previously described [11]. This vector was introduced into wild-type *Giardia* WBC6 as described above, using puromycin selection.

To make the DAP7268 quadruple knockout strain DAP7268quadKO, two *Giardia* antibiotic cassettes, BsdRgo and HygRgo were introduced sequentially into the Cas9DAP7268_gRNA908R expression strain above by electroporation of 20 to 40 ug of linear HDR template as previously described [11]. Each HDR template consisted of an antibiotic cassette flanked by the 750 bp of DAP7268 sequence immediately up- and downstream from the DSB site at position 908 (Supplemental Figure 1). The templates were synthesized by Twist Biosciences and cloned into their pTwist Amp High Copy vector. Linear HDR templates were amplified from this vector using Q5 DNA polymerase (NEB) and M13Forward and M13Reverse primers and purified using Zymo Research Clean and Concentrator-25 columns. Once knockout strains reached >50% confluence on full antibiotic selection, total DNA was extracted using the Zymo Research Quick DNA Miniprep Plus kit. Knock in of markers and loss of wild-type DAP7268 alleles was confirmed by PCR with primers 7268LeftF (5’-CTTATGCTGGCAGTGCGATCC-3’) and 7268RightR (5’-CTCTATAAGCTCCTCAACACTCTGC-3’), which bind roughly 800 bp from the DSB site, outside the homology arms. In contrast to our knockout of DAP16343[11], which required three markers, wild-type DAP7268 alleles were undetectable after knock-in of the second (HygRgo) cassette.

To make the DAP7268quadKO_DAP12139mNG strain, which stably expresses DAP12139 as a marker of the disc margin, we designed a fragment containing a DAP12139-mNeonGreen C terminal fusion, with DAP12139 expression driven by its native promoter (Supplemental Figure 2). The 2.9 kb fragment was synthesized (Twist Biosciences) and cloned into a unique MluI site that lies just upstream of the rpS28 promoter in the HygRgo cassette. The entire 6.3 kb DAP12139mNG_HygRgo HDR template was amplified, purified, and introduced via CRISPR knockout after the introduction of BsdRgo. The DAP7268quadKO_DAP12139mNG strain was confirmed by PCR as described above. For both strains, clonal cell lines were obtained by limiting dilution (see above).

### Live imaging of fluorescently tagged strains

The DAP12139 disc margin GFP fusion strain and DAP7268KO-DAP12139mNG strains were thawed from frozen stocks and cultured at least 24 hours prior to live imaging. Cells were iced, harvested, and resuspended in 1 ml warmed medium, and 300 µl aliquots were incubated in 96-well black glass bottom imaging plates (Cellvis, Mountain View, CA) or MatTek dishes for up to one hour at 37°C to promote attachment. Prior to imaging, the medium was replaced with warmed 1X HBS, and trophozoites were incubated under the same conditions for 30 minutes. Additional warmed 1X HBS washes were performed as needed during imaging to remove detached cells. DIC and epifluorescence imaging was performed with a Leica DMI6000B microscope as described previously [11]. Images were processed and disc contraction was quantified using kymographs using FIJI ^8^.

### Immunostaining and quantification of ventral disc dimensions in mutants

Prior to immunostaining, strains were grown to confluency and fixed in 12 ml tubes with a final concentration of 1% paraformaldehyde, pH 7.4 (37°C, 15 min). Detached cells were pelleted at 900 x g for five minutes. The pellet was washed three times with 2 ml PEM, pH 6.9 [12], incubated in 0.125M glycine (15 min, 25°C), and washed three more times with PEM. Fixed cell suspensions were then settled on to coverslips treated with 1mg/ml poly-L lysine for one hour. Coverslips were then washed 3X in PEM, and permeabilized with 0.1% Triton X-100/PEM for 10 minutes. Coverslips were blocked in PEMBALG for 30 minutes and incubated overnight at 4°C with primary antibodies for microtubules using anti-TAT1 (1:250, gift of K. Gull, University of Oxford) or ventral disc microribbons using anti-delta-giardin (1:100) or anti-beta-giardin (1:1000, gift of M. Jenkins, USDA) antibodies. Following the primary antibody incubation, coverslips were washed three times in PEMBALG and incubated with Alex Fluor 488 goat anti-rabbit and/or Alex Fluor 594 goat anti-mouse antibodies (1:1000; Life Technologies) for four hours at room temperature. Following incubation with the secondary antibodies, coverslips were washed three times each with PEMBALG and PEM and mounted in Prolong Diamond antifade reagent (Life Technologies). All imaging experiments were performed with three biologically independent samples.

Overall disc curvature, disc and bare area dimensions including perimeter and area, and surface contacts dimensions were quantified from > 300 immunostained images per condition using Fiji [15]. Disc surface contacts were quantified from CellMask-stained images.

### Super-resolution SIM 3D imaging of fixed cells

For super-resolution imaging of strains, three-dimensional stacks were captured with 0.125 µm intervals between slices. SIM microscopy used the Nikon N-SIM Structured Illumination Super-resolution Microscope with the 100X/NA 1.49 objective, 100 EX V-R diffraction grating, and an Andor iXon3 DU-897E EMCCD camera. The collected images were then processed using the “3D-SIM” mode, which involved reconstructing the images using three diffraction grating angles and three translations each. To evaluate the quality of the images, a reconstruction score was assigned to each image using the “Reconstruct Slice” mode. Only images with a score of 8 or higher were deemed acceptable for analysis. Additionally, both the raw and reconstructed images underwent further assessment using SIMcheck, a quality assessment tool developed by Ball et al. (. The final images are displayed either as maximum intensity projections or as 3D stacks, depending on the desired representation.

### Quantification of disc doming (angle of curvature)

To quantify the range of disc curvature, cells were attached to imaging dishes (MatTek) by incubating at 37°C for 30 min in 1X HBS, washed three times with 1X HBS to remove unattached cells, and resuspended in 100 µl of warm 1X HBS. 100 µl of warmed (37°C) 3% low melt agarose in 1X HBS was added to wells (final concentration of 1.5% agarose) to immobilize trophozoites for live imaging. To capture the entire disc, a Leica wide-field DMI6000B microscope was used to collect optical slices at 0.2 µm intervals for up to 8 µm. Disc doming or curvature images were generated by reslicing the stack laterally across the posterior portion of the bare area and angles were measured in ImageJ/Fiji [15]. For each experimental condition, at least 30 cells were measured on three separate days totaling 90 cells imaged and quantified.

### Quantification of shear forces of using a flow cell assay of attachment

To quantify their ability to resist shear forces in microscopic flow assays, the DAP7268KO strain, Cas9 vector control, DAP12139GFP, and wild-type trophozoites were initially incubated in Ibidi mSlide VI 0.4 flow chambers (Ibidi). Single focal plane images of attaching 1,000,000 cells were acquired using the μManager acquisition software [16] with the 10X objective using a Leica DMI 6000 wide-field inverted fluorescence microscope. Cells were allowed to attach for ten minutes prior to initiation of flow. Attached cells were counted by overlaying the pre-flow image over the post-flow image. The overall percent attached was calculated by quantifying the percentage of shear force resisting cells relative to the total number of cells.

## Organoid growth and TEER measurements

Human organoids were generated and maintained from duodenal biopsy specimens from healthy volunteers undergoing routine examinations at Charité-Universitätmedizin Berlin (ethics approval #EA4-015-13 by local authorities) were used to generate human organoids are previously as described [13]. Matrigel-coated (Corning, Tewksbury, NY) transwell cell culture inserts (0.6 cm2, 0.4-mm pores; Merck-Millipore, Burlington, MA) were then used to culture organoid derived monolayers (ODMs) using previously described methods [13]. For infections, wild-type and DAP7268KO trophozoites were passaged the day before infection experiments to guarantee logarithmic growth phase. Prior to infection, parasites were detached from culture tubes by incubation at 4°C for 20 minutes, pelleted and quantified using a Neubauer counting chamber. Parasites were then harvested by pelleting at 1000 x g for five minutes at °C and resuspended at the final desired concentration. The apical compartment medium of 8- to 10-day-old ODMs was replaced with complete TYI-S-33 the evening before infection to adapt cells to the TYI-S-33 medium. Medium was renewed before infection (i.e., TYI-S-33 in the apical and organoid differentiation medium in the basal compartment). *Giardia* trophozoites were added apically and incubated for up to 72 hours.

### Transepithelial electric resistance (TEER) measurements of epithelial barrier integrity

To assess the barrier integrity of disc mutants, transepithelial electric resistance (TEER) measurements were conducted. A Millicell ERS-2 Voltohmmeter (Merck-Millipore) equipped with an Ag/AgCl electrode (STX01; Merck-Millipore) was used for these measurements. The instrument was placed on a 37°C heating block to maintain a consistent temperature. To quantify the TEER values, the electric resistance of a blank transwell insert without cells was measured and subtracted from the raw resistance values obtained from the samples. Furthermore, the resulting values were standardized to account for a surface area of 1 cm2.

## Results

### Disc attachment models are defined by essential disc structural elements and flagellar movements

The ventral disc is a composite structure composed of an extensive MT spiral array and associated protein complexes including the MR-CB complex (body, Figure 1A). The disc also has a region of overlap, the overlap zone (oz, Figure 1A), that is key to its domed architecture. The disc margin marks the outside edge of the spiral MT array, including a gap at the point of the disc overlap zone (Figure 1A and [9]). The lateral crest is an additional repetitive structure is bound to the disc perimeter outside of the MT array, and has previously been argued to be contractile (Figure 1A, [3]). The ventral groove is a deformed region of the disc overlapping the ventral flagella, which were proposed in the hydrodynamic model to create a hydrodynamic current required for attachment by drawing fluid underneath an inflexible, non-contractile disc (Figure 1B).

Is the disc an inflexible, passive structure that merely maintains attachment forces generated by flagellar motility as assumed by Holberton, or is the disc an actively dynamic and flexible structure that independently mediates attachment forces? Overall disc contraction has been proposed to occur through shortening or compression of the crossbridges, reducing the spacing between the MTs of the disc spiral (e.g., disc compression, Figure 1C). This proposal implicates the MR-CB complex in an overall and evenly spaced compression of the disc during attachment, resulting in a lower volume underneath the disc and a smaller disc surface area and perimeter. Alternatively, the lateral crest has been proposed to constrict, resulting in an overall contraction of the disc. In this model, the spacing between the disc MT spiral may or may not be reduced. Lastly, the upper and lower layers of the disc could rotate or slide over each other, again resulting in an overall disc contraction. Spacing between disc MTs again may or may not be altered. Of course, the three models are not mutually exclusive.

### Direct evidence for disc contraction during trophozoite attachment to surfaces

To test the prediction that disc contraction occurs during parasite attachment to surfaces and to quantify disc contractile dynamics, we used quantitative time-lapse imaging to track disc movements in DAP12139GFP, which expresses a fluorescent tag marking the lateral crest. DAP12139GFP does not colocalize with primary components of the disc including the microribbon-crossbridge complex (Figure 2A and [3]). Three-dimensional line scans were drawn at three regions of the disc margin including the overlap zone (OZ), ventral groove (VG), and the disc body, to define disc margin movements with kymograph analyses (Figure 2B,C,D and Supplemental Figures S1-S3) corresponding to disc contractile movements associated with overlap zone rotation (Figure 2B), disc compression (Figure 2C) and disc constriction (Figure 2D). First, we observed that the overlap zone region rotates clockwise toward the disc center (Figure 2B, asterisk and Supplemental Figure S1). Second, by measuring changes in the horizontal distance between the disc edges (Figure 2C), we observed disc compression (Figure 2D and Supplemental Figure S2) that averaged 0.3-0.5 µm during attachment but ranged up to about 0.8 µm (Figure 3E). Lastly, lateral crest constriction also occurred during attachment (Figure 2F,G,H and Supplemental Figure S3), mirroring disc overlap zone rotation with clockwise constriction of the disc margin toward the ventral groove and causing the opening of the disc margin to increase on average about 0.4 µm (Figure 2G,H). In total, we quantify disc contraction of about 0.5 µm on average during attachment that is attributable to the three types of quantifiable movements (Figure 2I) that occur within 15 seconds (overlap zone rotation) to 30 seconds (disc compression and constriction).

**Figure 2:**
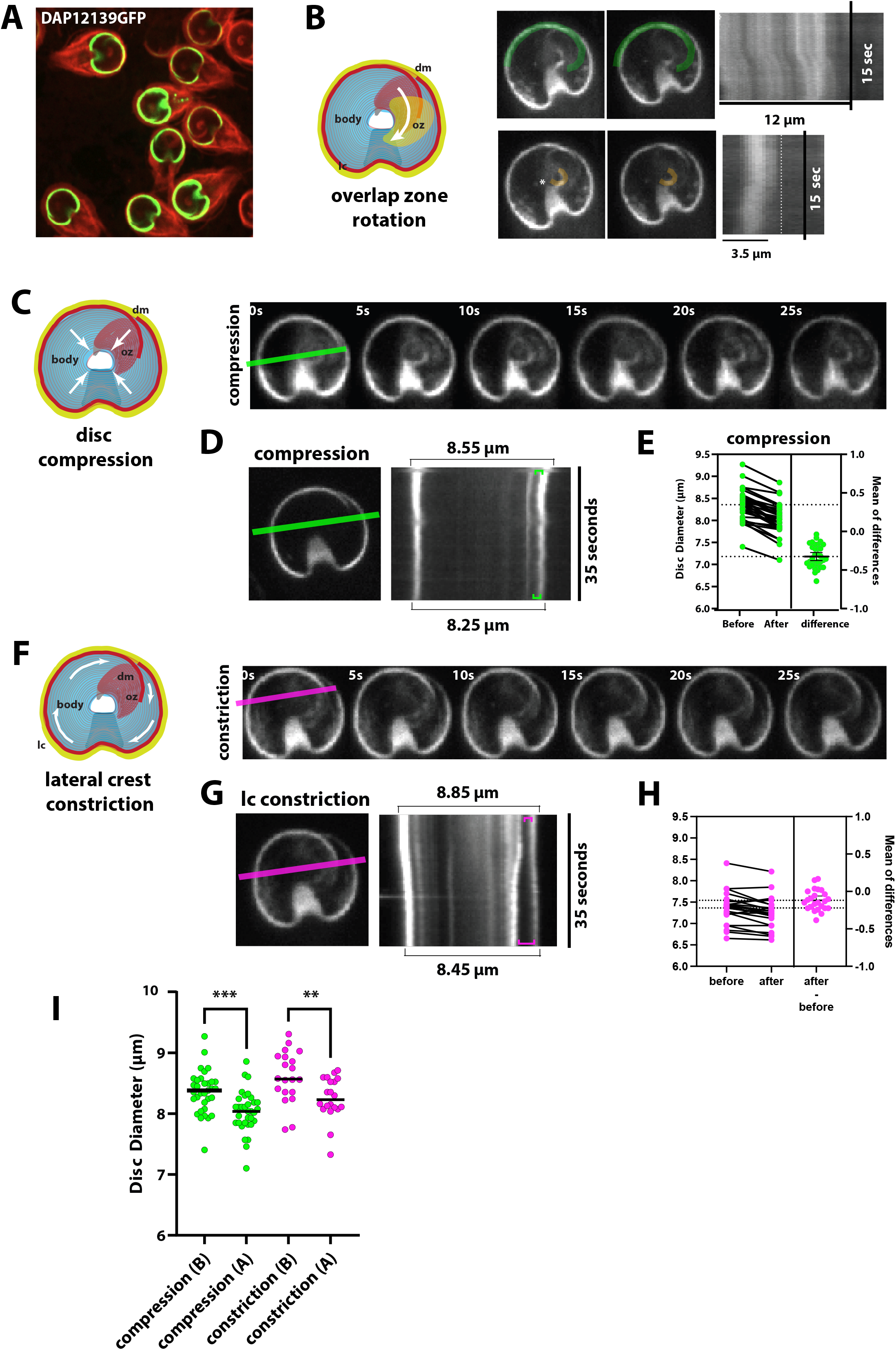
Three types of disc movements contribute to overall disc contraction. Using a lateral crest marker to quantify disc movements (panel A, DAP12139GFP), disc movements were quantified using live imaging of over 20 cells with kymographic analyses. In B, the rotational movements of the overlap zone are defined and quantified. In C, overall disc compression movements (green line) that occur within 25 seconds in attaching trophozoites are quantified using kymography analysis (D) with summaries of movements of over 15 cells (E). In F, movements leading to constriction of the lateral crest (magenta line) resulting in disc contraction are quantified in kymographs (G) and with over 15 cells summarized (H). In panel I, disc contraction measurementss indicate contraction of up 5-10% of disc diameter.

**Figure 3:**
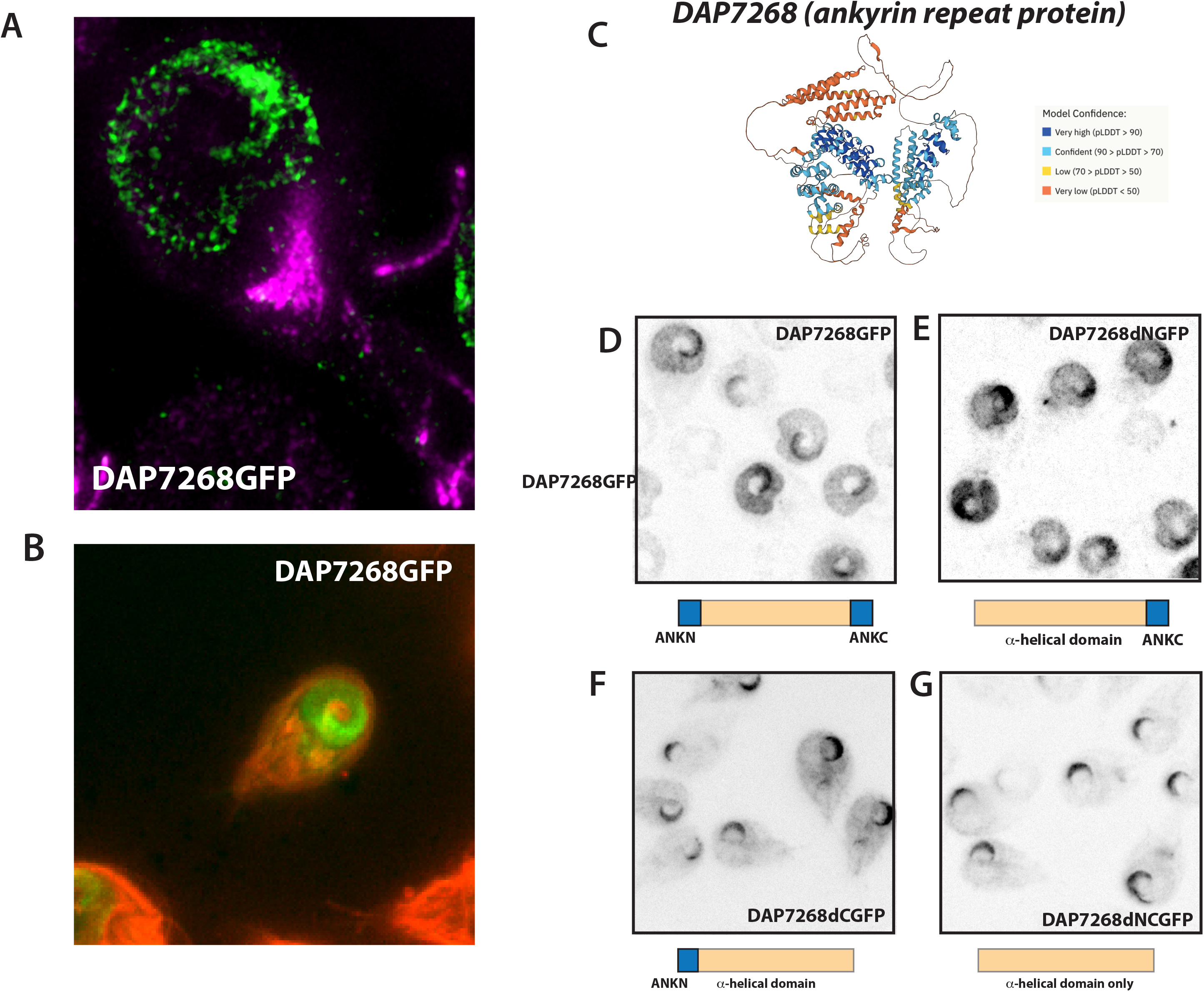
Both N- and C-terminal ankyrin repeat domains are necessary for localization of DAP7268 to the disc body and overlap zone. Structured illumination (SIM) imaging (A) and wide field imaging mNeonGreen-tagged DAP7268mNG (green) highlights the localization of this DAP to the disc overlap zone and disc body relative to the MT cytoskeleton (magenta) and to membrane-stained cells (B, CellMask, red). In C, an AlphaFold structural predictions shows high confidence with the N- and C-terminal ankyrin repeats (ANKN or ANKC), but less confidence with other alpha helical regions of DAP7268 (C). In panels D-G, localization of various DAP7268mNG constructs with N-, C-, or both ankyrin repeat regions are indicated in inverted false colored images (full or partial DAP7268 GFP fusions = black).

### Interrogating the role of the disc overlap zone ankyrin repeat protein DAP7268 in disc contractility

Using disc-based proteomics with subsequent protein tagging, we have previously shown that DAP7268 is a disc-associated protein that localizes to the overlap zone and disc body (Figure 3A-B). DAP7268 also has one N-terminal and one C-terminal ankyrin repeat domain (Figure 3C), like 30 other DAPs [3]. A predicted alpha helical domain lies between the terminal ankyrin repeat domains. To determine the localization role of the ankyrin repeat domains, we expressed truncated forms of DAP7268 fused to a C-terminal GFP. This ankyrin domain analysis (Figure 3D,E,F,G) indicates that the N terminal ankyrin domain is necessary for localization of DAP7268 to the disc overlap zone (Figure 3E, no oz localization). In contrast, the C-terminal ankyrin domain is necessary for localization to the disc body (Figure 3F). Deletion of both ankyrin repeats has a similar localization as just the C-terminal ankyrin.

### DAP7268 quad knockouts have a “Baumkuchen” disc phenotype defined by multiple overlapping disc layers

In initial DAP morpholino-based transient knockdown screens, we found that the DAP7268 KD strain had aberrant disc structure (Figure 4C,D). Using our lab’s new methods to create quadruple allele deletion mutants (“quad knockouts”, Methods), we confirmed a quad knockout using blasticidin (B) and hygromycin (H) repair templates. Confirmation through genomic sequencing indicated that two alleles were disrupted by the B marker, and two by the H marker (Supplemental Figures 4 and 5).

**Figure 4.**
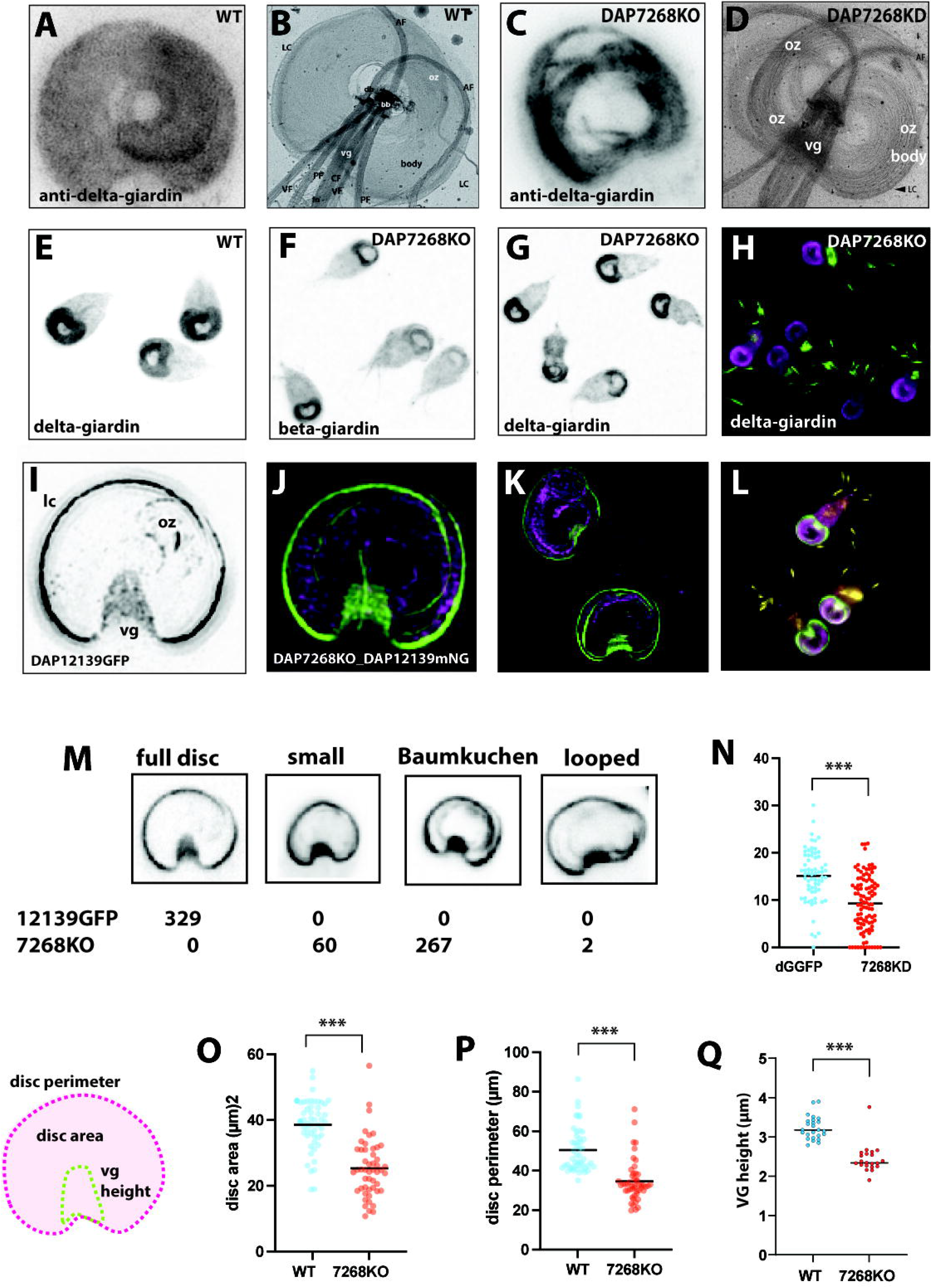
Quadruple allele DAP7268KO mutants have aberrant multi-layered, yet smaller discs with multiple lateral crests. SIM imaging of DAP7268KO and wild-type discs immunostained with delta-giardin or beta-giardin highlight aberrant discs. The characteristic spiral array of wild type discs is shown in (A,B) seen with SIM with delta giardin immunostaining (A) and negative staining of extracted disc (B). In C, the aberrant structure of the DAP7268KO is (delta giardin immunostaining to mark disc microribbons). In (D), the similar aberrant structure is seen in negative staining of the extracted disc in the morpholino-based knockdown DAP7268KD. The 100% penetrance of the DAP7268KO phenotype is seen in immunostained beta- and delta-giardin images (E-G) and with anti-delta giardin (magenta) and anti-alpha tubulin (green) immunostaining (H). To visualize the lateral crest in the DAP7268KO mutant, the DAP12139mNG lateral crest marker (green) was integrated into DAP7268KO strains enabling the visualization of multiple lateral crests in the DAP7268KO mutant background. Discs were immunostained with delta-giardin (magenta, J, K) and microtubules were also immunostained with anti-alpha tubulin (TAT1) antibodies (yellow, L). In M, DAP7268KO mutant phenotypes were scored based on the three most common mutant phenotypes (small, Baumkuchen, or looped). Quantitation of overall disc doming angle of curvature (N), disc body area (O), disc perimeter (P), and ventral groove height (Q) are summarized for delta-giardin GFP strains (dGGFP), DAP7268 morpholino knockdown strains (7268KD), 7268KO, and wild type strains.

**Figure 5:**
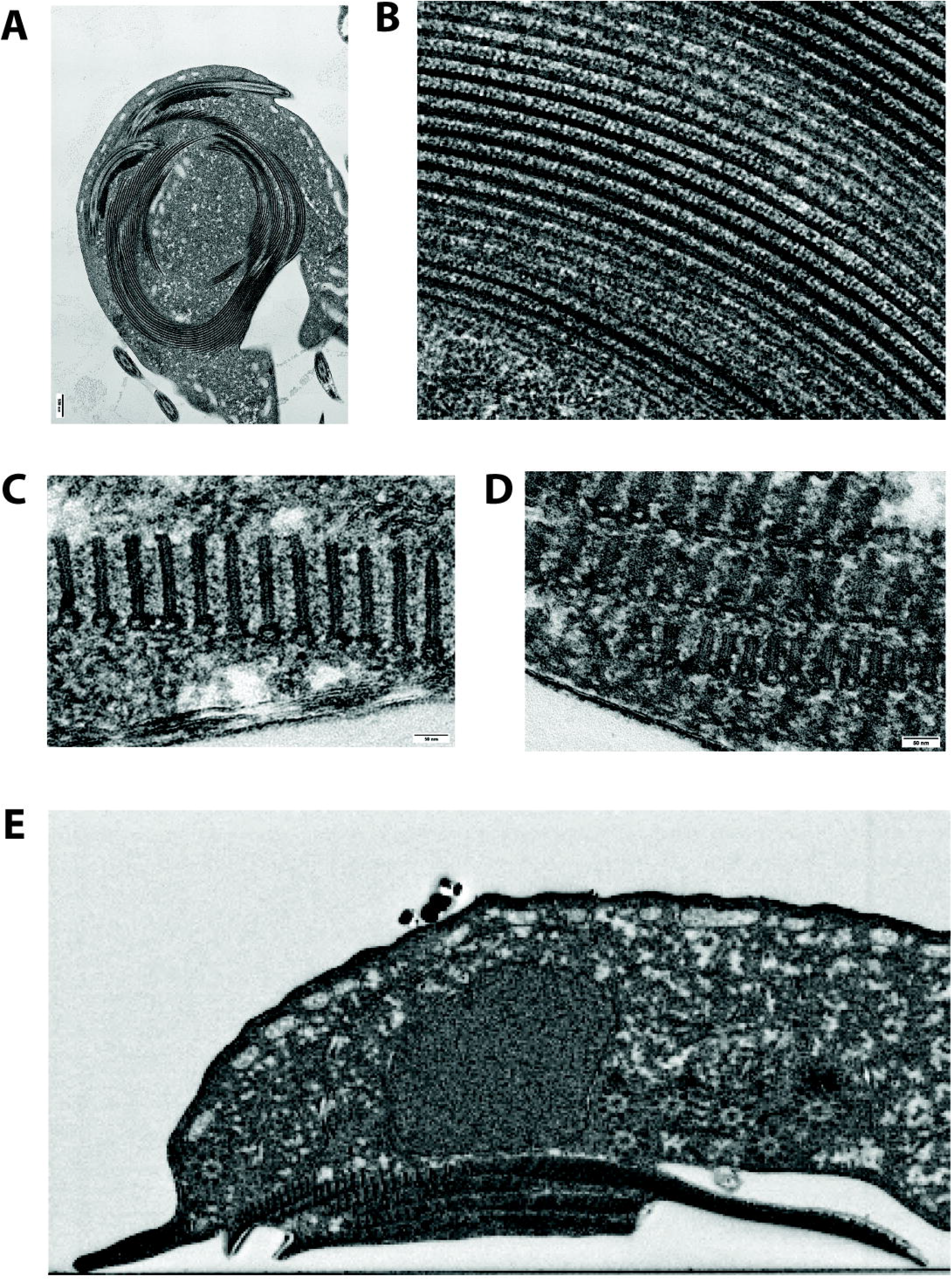
DAP7268KO mutants are defined by a multilayered “Baumkuchen” phenotype with multiple aberrant MR-CB complexes and multiple lateral crests. Transmission electron microscopy with sectioning DAP7268KO whole cells highlight the aberrant and extra spiral MT arrays (MTs) (A). In mutants, the microribbon-crossbridge complexes have aberrant spacing (B). DAP7268KO strains have multiple overlap zone layers as seen in the comparison of TEM cross sections of wild-type MR-CB complexes (C) with extra layered and shortened MR-CB complexes (D). Panel E shows the additional overlap zone layers, each with MR-CB complexes stacked above one another in serial block face SEM sections.

Immunostaining of the DAP7268quadKO strain using anti-delta Giardia antibodies that mark disc MT-CB complexes indicated that the disc structure was severely disrupted (SIM, Figure 4C; negative staining, 4D) as compared to the wild-type control strain. To visualize the lateral crest region in DAP7268 quad knockouts, we also created an additional strain (DAP7268quadKO_DAP12139mNG) that included a mNeonGreen (mNG) C-terminally tagged lateral crest marker (DAP12139, Figure 4) adjacent to the hygromycin cassette, and confirmed this insertion allele using long read sequencing (Supplemental Figure 6). By immunostaining for both the microtubules (anti-TAT1) and the MT-CB complex of the disc (anti-delta-giardin and anti-beta-giardin), we discovered that DAP7268 quad KOs had an aberrant disc with disc multiple layers as well as a large central region (bare area). The extra layers are readily observable as extra disc margins (marked by DAP12139mNG). DAP7268KO mutant disc phenotypes are 100% penetrant (Figure 3D). We termed the layered disc phenotype “Baumkuchen” after the traditional multilayered German cake of the same name (Figure 3D). Baumkuchen phenotypes are the most penetrant of DAP7268KO disc defects, as compared to “small” discs or discs with extra partial spirals (“looped”). Overall, the area of DAP7268KO discs was smaller (Figure 4O) and the discs were more flattened (Figure 4N). DAP7268KD discs are also much less domed that wild type (Figure 4N). Furthermore, DAP7268KO mutant discs are on average about 80% smaller with respect to disc perimeter, disc area and the height of the ventral groove (Figure 4O-Q)

Knocking in DAP12139mNG in the context of the DAP7268 quad KO allowed us to quantify aberrant lateral crests in this DAP mutant. We observed multiple lateral crest regions (Figure 4) that in some cases bulged out into additional aberrant lateral crests. Using TEM, we confirmed that the aberrant disc structure of the Baumkuchen phenotype included additional layers (Figure 5). We observed upwards of four or five overlapping layers in some cases, each with MTs and MR-CB complexes. In addition, each extra layer had aberrant microribbons that were of different heights and irregular spacing (Figure 5). We confirmed the duplication of the lateral crest using extracted cytoskeletons of the DAP7268 morpholino knockdown visualized with negative staining. In extracted cytoskeletons, the extra discs layers were again visible, with additional overlap zones and duplicated lateral crest structures causing a more circular, doubled disc (Supplemental Figure S7, arrows).

### DAP7268KO Baumkuchen mutants lack disc contractility

The presence of multiple or aberrant disc edges/lateral crests in Baumkuchen mutants implied that mutants could also have defect in disc edge contractility as demonstrated in Figure 2. Imaging attaching DAP7268KO_DAP12139mNG strain mutants allowed us to quantify disc contractile dynamics in DA7268KO mutants using their integrated DAP12139mNG the lateral crest marker (Figure 6). As seen in live imaging (Figure 6A and Supplemental Figures S8-S11) and associated kymographs (Figure 6B), Baumkuchen mutants have significantly decreased overall disc contraction. With respect to overlap zone rotation, mutants had slight rotation in the opposite direction as we observed in DAP12139GFP strains (Figure 2). With respect to both disc compression and constriction measurements, we determined DAP7268KO mutants have on average only 0.2 µm contraction as compared to wild-type strains (Figure 6C,D).

**Figure 6:**
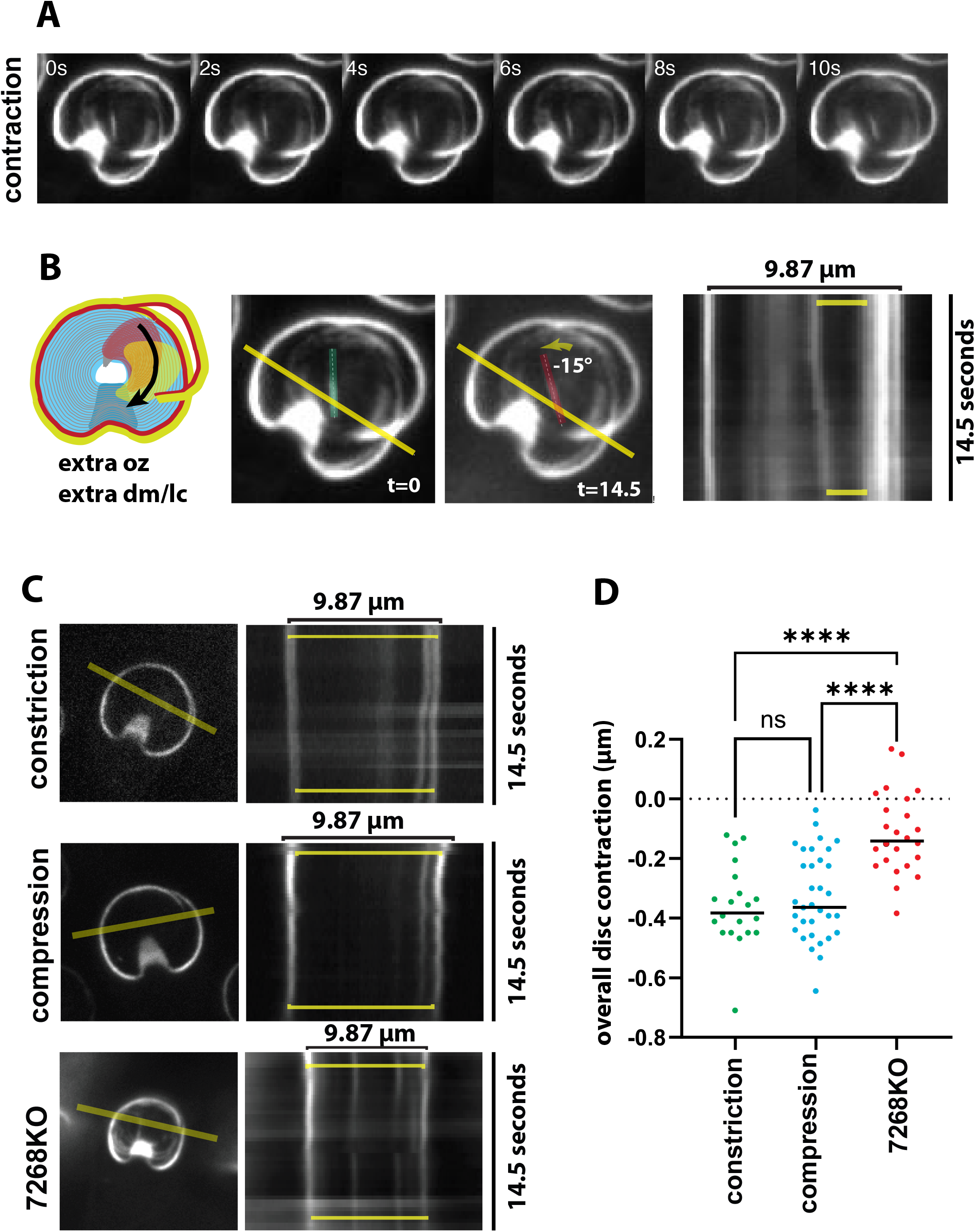
DAP7268KO “Baumkuchen” mutants lack proper disc contractility. To test disc contractility or other disc movements the Baumkuchen mutant, we integrated the DAP12139mNG marker into the DAP7268KO strain and tracked disc contraction over several seconds (A). The schematic in B indicated the doubled and aberrant lateral crests that showed reduced and backwards overlap zone rotation movements as demonstrated using live imaging and kymograph analysis (B). Baumkuchen mutants also have markedly reduced disc constriction or compression (C, D).

### Baumkuchen mutants have decreased attachment surface contacts and resistance to shear forces

Membrane contacts are diagnostic of the level of cellular attachment to surfaces during early attachment “stages” [12]. Initial cellular contacts in Stages I and II include the ventrolateral flange and disc edge, and later Stage III and IV contacts include the “tail” regions and the lateral shields and the bare area in the center of the disc. After staining with the membrane dye CellMask Orange, we determined that membrane contacts with the surface were altered in the DAP7268KO mutant. We observed larger membrane regions of the bare area contacting the surface (Stage IV+ba), and in some cases the bare area membrane lacked any contacts with the surface (Figure 7A). In comparison to wild-type surface contacts which are primarily at stage IV (including lateral crest, lateral shield, and bare area contacts), the majority of DAP7268KO mutants reached a modified stage IV stage which lacked surface contacts associated with the bare area (Stage IV-LC). We also noted in light and electron microscopy that the central disc region (bare area) was larger in mutants, and this is reflected in the size of the bare area surface contacts observed (Figure 7A, Stage IV+B).

**Figure 7:**
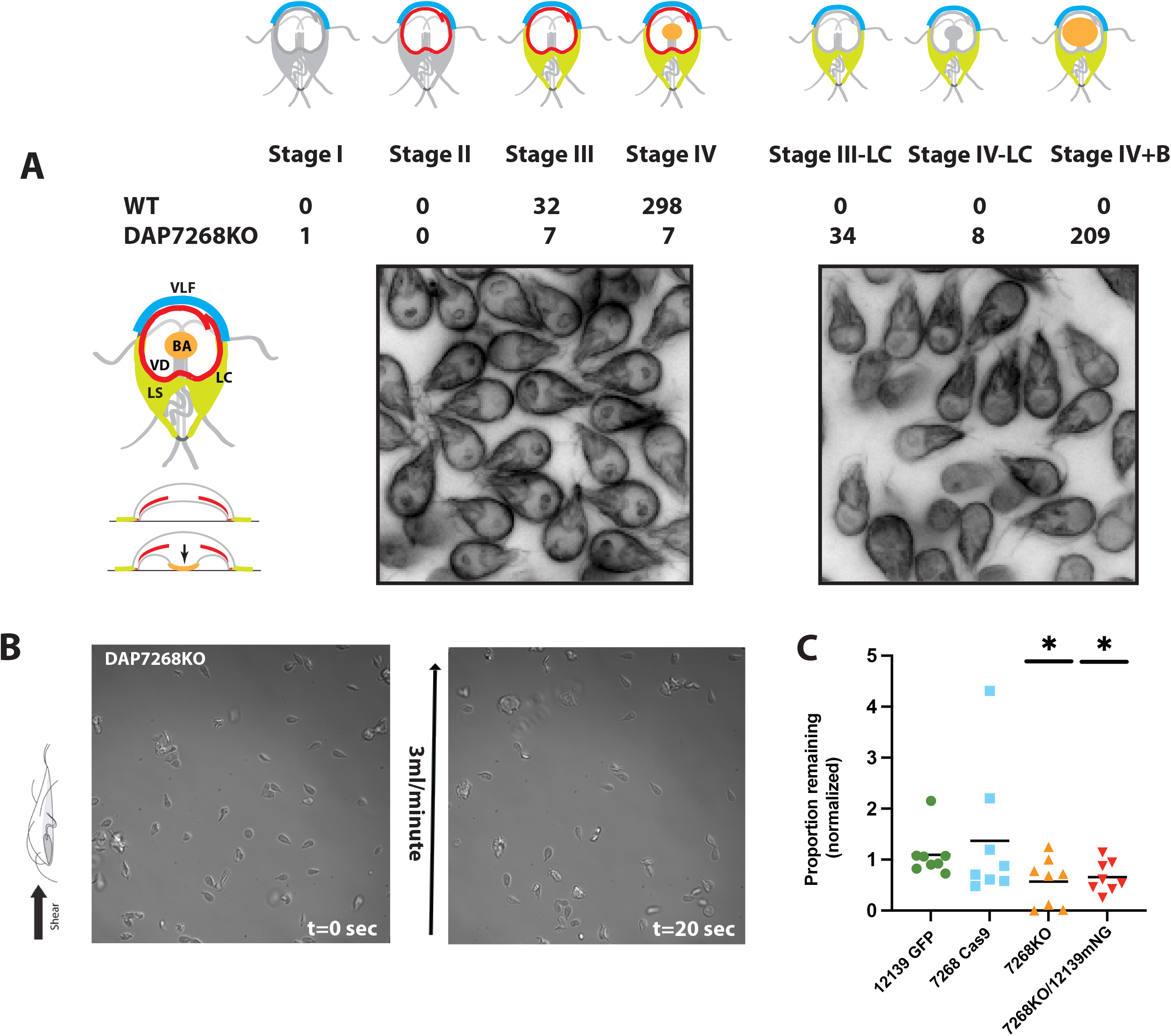
DAP7268KO strains have significantly reduced surface contacts and reduced ability to resist flow stress associated with proper attachment. In panel A, surface contacts associated with various early stages of trophozoite attachment (stages I-IV, [12]) are scored in both DAP7268KO and wild-type strains. DAP7268KO mutants lack canonical surface contacts, and instead have three additional attachment stage categories: stage III-LC and stage IV-LC which both lack lateral crest contacts, and stage IV+B which is defined by aberrant bare area contacts connecting to the lateral crest. Shear stress assays using 3 ml/minute flow (B) highlight the severe defects in the ability of DAP7268KO mutants to resist shear stress, specifically, the reduced proportion of attached cells following 20 seconds of shear stress as compared to the 7268Cas9 plasmid alone strain, wild-type, DAP12139GFP tagged, and DAP7268KO/DAP12139mNG strains (C). N = 7 to 8 cells. Asterisks indicate p values < 0.01 in t-tests.

To evaluate shear forces of attachment, we used a novel quantitative shear stress assay that uses time lapse imaging of trophozoites in commercial flow chambers (Methods). Wild-type, the Baumkuchen quad KO, DAP12139GFP and the DAP7268KO_DAP12139mNG strains were flowed into chambers, allowed to attach, and then challenged with a range of shear forces. Mutants and tagged strains were normalized to wild-type. Less than 40 percent of the attached parasites resisted 3ml/min of flow (∼4dyn/cm2) Baumkuchen DAP7268KO and DAP7268KO/DAP12139mNG strains trophozoites had significant decreases in their ability to resist 3ml/min of flow (Figure 7B,C).

### Significantly reduced epithelial barrier breakdown in Baumkuchen mutants using human intestinal organoid derived monolayers

2D human intestinal organoid-derived monolayers (ODMs) undergo maturation over a period of approximately 9 days in culture. During this time, the ODMs reach a height of around 20 μm and exhibit the characteristic polarized shape of enterocytes, including the presence of microvilli and well-developed cellular junctions [36]. As the ODMs mature, there is a gradual increase in trans-epithelial electrical resistance (TEER), which stabilizes at approximately 250 Ω · cm2 after 9 days of culture. TEER serves as a measure of the integrity of the barrier function, and its decrease is observed in *Giardia* infections of ODMs, indicating disruption of the barrier function [36].

To investigate the impact of wild-type and Baumkuchen mutant *Giardia* trophozoites on the degradation of barrier function, ODMs were infected with both wild-type trophozoites and two DAP7268KO clones (7268-B2 and 7268-C7) or treated with vehicle in mock infections (no *Giardia*) at different multiplicities of infection (MOI) ranging from 1 to 10. In wild-type infections, the normalized TEER exhibited a dose- and time-dependent decrease after 24 hours of infection, reaching its peak at 48-54 hours (Figure 8A, B). In contrast, both DAP7268KO clones showed significantly higher normalized TEER compared to the wild-type control infections.

**Figure 8:**
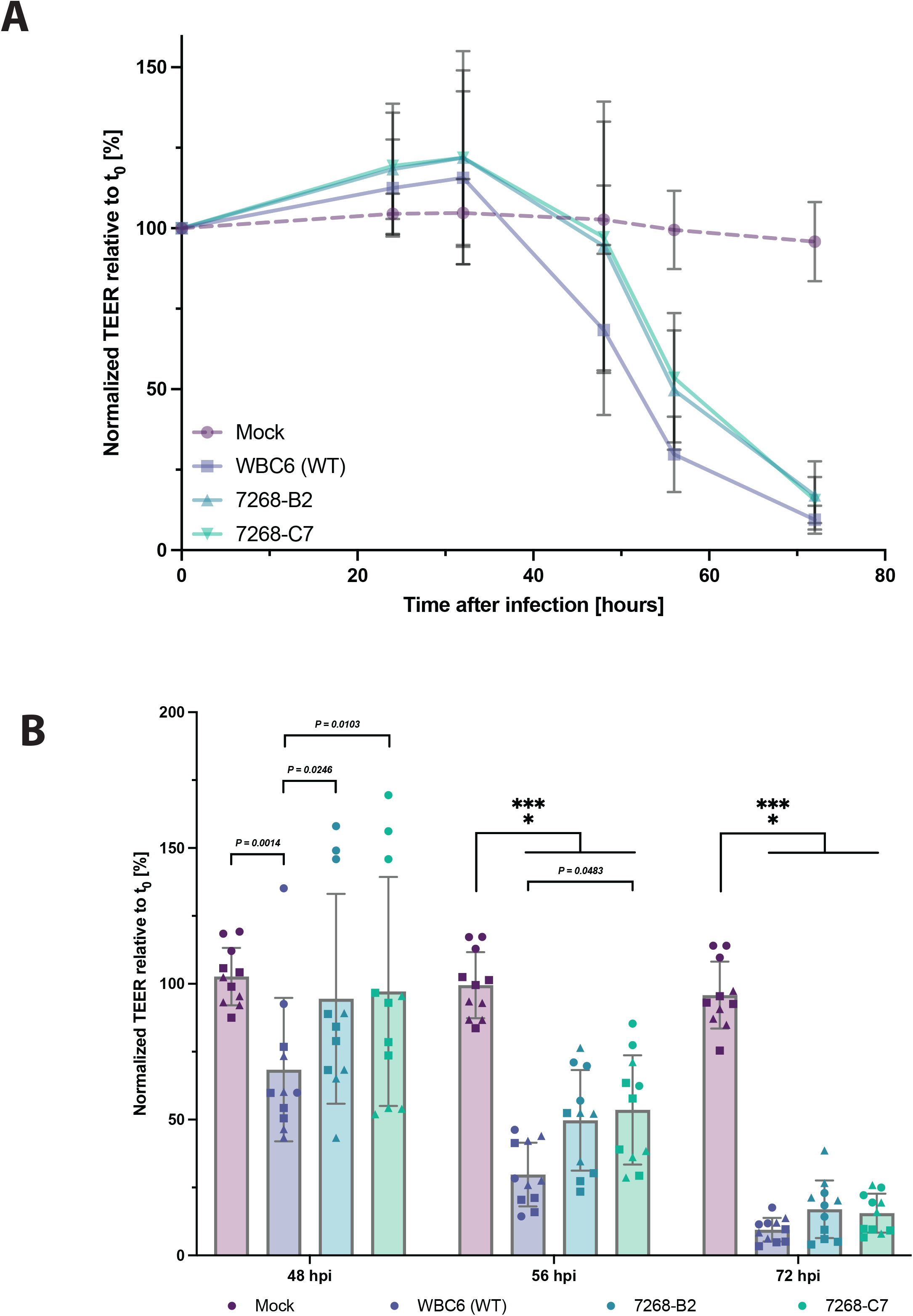
DAP7268 attachment mutants have a significantly reduced impact on enterocyte barrier degradation. TEER measurements of ODM-based infections of wild-type (WBC6) trophozoites and two DAP7268KO disc mutant clones (DAP7268-B2 and DAP7268-C7) are compared with mock (no *Giardia*) infections in an 80-hour time period (A). In B, relative TEER measurements are compared at representative time points (0, 48, 56, and 72 hours post infection (hpi)), with significant differences indicated by T-tests (***).

## Discussion

*Giardia*’s attachment via the ventral disc to either the host epithelium or to inert surfaces is defined by its rapid, reversible, and dynamic nature, enabling the parasite to resist peristaltic flow in the gastrointestinal tract and colonize the host. Numerous hypotheses have been proposed over the past 50 years to explain attachment mechanisms involving the ventral disc and/or lateral crest. These hypotheses encompass hydrodynamic suction forces generated by flagellar motility [5,8,17], cytoskeletal-based contraction of the disc, lateral crest, or ventrolateral flange [7,18–20], microtubule sliding dynamics [21], and lectin-based [22–25] or osmolarity-based mechanisms [26]. Among these, the “hydrodynamic suction model” has gained prominence based on early observations of *Giardia muris* attached to glass slides and modelling of fluid dynamics modeling [8].

An early assertion of the hydrodynamic suction model is that the ventral disc serves as a rigid, non-contractile structure, yet other early observations suggested a more contractile disc and lateral crest that engages and directly grips or “clutches” the host epithelium [7]. This disc contractility has been proposed to account for the formation of disc-shaped tissue damage or “craters” when trophozoites attach to microvilli. Such conflicting observations and insufficient empirical evidence supporting attachment mechanisms such as flagellar-based hydrodynamic suction have led to incongruent attachment models, however [5,7,8,17–25]. Moreover, the lack of advanced genetic tools for creating stable attachment mutants and a lack of robust *in vitro* and *in vivo* infection models has hindered investigations into whether contractility-based attachment directly contributes to the observed host cellular damage.

To gain insights into disc attachment mechanisms, we examined the contractility of the disc and lateral crest. Through live imaging (Figure 2) and analysis of a DAP contractility mutant with what we term the “Baumkuchen” phenotype (Figures 3-7), we provide direct evidence of disc and lateral crest contractility. While the specific molecular mechanisms underlying these contractile movements are yet to be determined, our results are consistent with earlier models put forth by Feely et al. [7] and Narcisi et al. [18], yet in the absence of actin-based contractility. This discovery of disc movements associated with contractility is pivotal for advancing our understanding of *Giardia* attachment mechanisms and their impacts on the host epithelium.

### Three types of dynamic movements of the ventral disc during attachment

We define and quantify three types of disc dynamic movements that occur in seconds in attaching cells and result in overall disc contraction: “**disc compression**”, “**lateral crest constriction**”, and “**overlap zone rotation**” (Figure 1 and Figure 2). Each of these movements are not mutually exclusive, and all may contribute to overall disc contraction we report here. Overall disc contraction in attaching cells reduces the diameter and the volume of discs by roughly 10-15 % (Figure 2). Changes in the relative positions of various disc components, such as MR-CB complexes, the overlap zone region, and the lateral crest have the potential to either expand or contract the diameter of the disc resulting in the characteristic “gripping” of the host microvilli by the ventral disc [7]. Although indirect mechanisms involving anterior flagellar movements may also contribute to disc contractility, there is currently no supporting evidence for this notion.

The first mode of disc contractile movement, which we term “**disc compression**,” is expected to act uniformly throughout the disc. A potential mechanism could involve the compression of crossbridges connecting adjacent microtubules (MTs) while maintaining a 50-nanometer spacing in the disc MT spiral. Dynamic movements of microribbons, composed of SF-assemblins (beta-giardin, delta-giardin, SALP1 [3], could also contribute to overall disc compression. Although Holberton [5] initially proposed this hypothesis regarding disc MR-CB complex compressions, it remains unconfirmed.

The type of second contractile movement we observe involves the direct “**constriction**” of the lateral crest or disc margin area of the disc (Figure 1C). Early observations of attaching trophozoites proposed that contractile activity in the lateral crest leads to the compression of the brush border microvilli [27](Erlandsen and Chase 1974, Takano and Yardley 1967, Balazs and Szatloczky 1978). These contractile forces were initially attributed actin and actin-associated proteins including myosin-based immunostaining of actin-binding proteins using heterologous antibodies (Feely 1982). However, later genome sequencing revealed that these localizations were artifacts, as only actin itself and no canonical actin binding proteins are present in *Giardia*. Furthermore, *Giardia* actin does not directly localize to the disc itself, but may interact with the disc through actin-associated disc proteins or DAAPs [28]. Alternatively, the constriction of the lateral crest could reduce the disc perimeter, and the mechanisms of constriction could be governed by specific lateral crest protein complexes or DAPs. Our laboratory has identified 30 disc-associated proteins (DAPs) that localize to the lateral crest and disc margin including ankyrin repeat proteins that may contribute to the lateral crest constriction that ultimately results in disc contraction. The ankyrin repeat protein DAP12139 used to track contractile movements (Figures 2 and 7) is likely associated with the lateral crest as the GFP staining does not colocalize with disc components including microribbons (Figure 2).

The third dynamic disc movement involves the rotation, or possible sliding, of the upper and lower layers of the overlap zone, a movement we term “overlap zone rotation” (Figure 1C). The physical connection between the upper and lower layers is not well understood, and the mechanism by which overlap zone DAPs or other DAPs may mediate these movements remains unclear. Additionally, these disc movements could be associated with the constriction of the lateral crest.

### Phenotypic analyses of DAP7268KO Baumkuchen mutants support an evidence-based model of attachment that requires disc and lateral crest contractility

The game-changing new genetic methods used here to create the stable, isogenic DAP7268 quadruple deletion mutant alleviate issues of mutant phenotypic transience [29] or incomplete penetrance [14] that can confound and limit both *in vitro* and *in vivo* assays of attachment. Because plasmids, such as CRISPRi plasmids, are readily lost off positive selection [14], DAP knockouts are also critical for *in vivo* analyses of *Giardia*-host interactions and pathology.

Initially identified in DAP morpholino screens, DAP7268 knockdown and knock out mutants of result in gross structural defects of the disc spiral including duplication or additional “layering” of the disc and lateral crest (Figure 4). We have coined the term “Baumkuchen” to describe this mutant disc phenotype, as it resembles a traditional German cake with multiple thin layers that resemble tree rings. DAP7268 consists of one N-terminal and one C-terminal ankyrin repeat domain linked by a central alpha-helical yet non-homologous region (Figure 3) and is among more than 30 ankyrin repeat containing disc-associated proteins (DAPs). DAP7268 is also one of the 31 DAPs that specifically localize to the “overlap zone,” and is specifically positioned at the outer edge of the overlap zone (Figure 1). Ankyrin repeats are generally recognized for their involvement in protein-protein interactions (Li et al., 2006), and some ankyrin proteins have been shown to directly interact with or bind to microtubules (Bennett and Davis, 1981; Davis and Bennett, 1984). Furthermore, in the apicomplexan protist *Toxoplasma gondii*, ankyrin repeat proteins link structural and motor proteins with the tubulin-rich conoid fibers (Long et al., 2017). In *Giardia*, our previous genetic analysis of another disc associated ankyrin-repeat protein DAP5188, which localizes to the entire disc and overlap zone, revealed its role in limiting interphase disc edge MT dynamics and promoting disc hyperstability [10].

DAP7268 knockout mutants also have irregular microribbons with uneven heights and spacing (Figures 4 and 5), disrupting the microribbon-crossbridge (MR-CB) complexes crucial for the overall domed architecture of the disc [30]. The disc spiral MT array’s edge is defined by the “disc margin” and is surrounded by the “lateral crest,” which makes contact with the ventral surface and has a gap at the overlap zone ([3] and Figure 1). Depletion of DAP7268 leads additional disc layers and additional lateral crests (Figures 4 and 5). The Baumkuchen phenotype supports a negative regulatory role for DAP7268 has in disc assembly, preventing additional layers of the disc MT spiral and lateral crest from forming during cell division. The presence of duplicated lateral crest regions, marked by DAP12139mNG in both wild-type and DAP7268 knockout mutants, further supports the idea that the lateral crest assembles after the disc MT array (Figures 4 and 5).

DAP7268KO Baumkuchen mutants also have significantly reduced (about 60% less) disc contraction and lack overlap zone rotation or other contractile movements (Figure 6). The lack of disc contractility in mutants is likely due to aberrant and extra disc edges/lateral crests. Disruption of the MR-CB complex in the DAP7268KO is another mechanism that may contribute to the observed lack of disc contractility and attachment (Figures 6 and 7). The lack of disc contractility in Baumkuchen mutants importantly results in decreased ability of parasites to resist shear forces in flow chambers (Figure 7) or create proper surface membrane contacts that require e a lateral crest seal [9]. During the early stages of attachment, trophozoites make initial contact with the surface using a specialized anterior structure called the ventrolateral flange (VLF), as first described by Holberton [5]and later confirmed by Erlandsen [19]. Recent studies have further supported the existence of focal contacts initiated by the VLF in the adhesion of parasites on laboratory surfaces [20]. These focal contacts contribute to the formation of a seal, which plays a crucial role in maintaining negative pressure underneath the disc, independent of flagellar beating. The presence of this seal marks the transition from “attaching” trophozoites to “attached” trophozoites (Figure 7 and [9]). The lateral crest seal serves two main purposes: it firmly secures the trophozoite to the attachment surface and restricts the flow of fluid entering or exiting from beneath the disc on smooth, non-elastic surfaces such as slides. Additionally, this seal positions the dorsal ventral disc membrane and peripheral vacuoles in direct contact with the microvilli of the host’s brush border.

Thus, we contend that disc contraction dynamics are necessary and likely directly responsible for the force generation required for attachment. We propose a “**disc contraction**” model of attachment that incorporates overall disc contraction with subsequent constriction of the lateral crest to create a closed seal around the disc perimeter. Such dynamic movements are necessary for attachment and resistance to hydrodynamic flow on inert surfaces, and likely result in “gripping” or “grasping” of parasites on host enterocytes.

### Disc contraction and attachment are necessary for intestinal barrier function breakdown associated with giardiasis

Direct cellular damage caused by extracellular attachment via the ventral disc has been assumed for decades, but not directly proven. Early reports also suggested that upward curvature of the brush border membrane of epithelial cells seen underneath the ventral disc in attached trophozoites was caused by attachment suction-based forces. *Giardia* infections are known to either directly or indirectly cause various types of damage to the host epithelium. Attached trophozoites could directly cause observed epithelial damage including reduced barrier function and integrity, to microvilli damage and shortening, degradation of the mucus layer, decreased nutrient absorption, and apoptosis. The direct or indirect impact of *Giardia’s* extracellular attachment on the observed cellular pathology in *Giardia* infections has been largely overlooked in previous investigations of the disease’s mechanisms.

The recent development of primary organoids, derived from small intestinal stem cells [6, 36] has opened up new opportunities for understanding *Giardia* pathogenesis. 3D cultures, derived from human duodenal biopsy specimens, are used to create 2-dimensional transwell systems known as organoid-derived monolayers (ODMs). Infections of ODMs with *Giardia* cause a gradual disruption of the epithelial barrier, directly correlated with the duration of infection and the parasite load. Infections trigger changes in gene expression related to ion transport and the structure of tight junctions in ODMs. Recent studies using ODM models have also revealed alternative mechanisms by which *Giardia* induces cellular pathologies. Inhibiting the activity of cysteine proteases and caspases, which are enzymes suggested to be virulence factors in giardiasis, does not prevent the breakdown of the epithelial barrier as measured using trans-epithelial electrical resistance (TEER) or prevent the occurrence of apoptosis. Further, we recently showed that attachment is required for complete barrier breakdown. Infections with the disc MBPquadKO mutant with significant structural and attachment defects, have markedly reduced impacts on barrier integrity as compared to wild-type *Giardia* infections of ODMs.

Here we provide additional support for the direct link between attachment and intestinal barrier breakdown. Using infections with the 100% penetrant Baumkuchen mutants that lack disc contract contractility, we again see a similar reduction the degree of barrier breakdown in mutants as compared to wild type infections (Figure 8). Both DAP7268KO clones show significantly limited ability to induce barrier breakdowns, which highlights the crucial role of parasite attachment in causing complete barrier breakdown in this ODM model of giardiasis. Other factors are likely still involved since we do not observe a complete absence of barrier breakdown, but the close and intimate association provided by extracellular contractility-based attachment is necessary for host cellular pathology and likely contributes to the systemic symptoms observed in giardiasis.

### Conclusions

The modern molecular genetic and live imaging approaches used in this work bridge the gap between early observations of disc contraction and the presumed effects of attachment on the host epithelium. While live imaging of movements of the disc resulting in disc contraction on inert surfaces would likely result in a gripping or grasping behavior of parasites on host enterocytes, the precise mechanism underlying these movements remains unknown. DAPs comprising specific structural elements of the disc may generate the forces required for disc contraction, yet these also remain to be discovered. Future investigations should aim to identify the specific DAPs involved in the contractile mechanisms of the disc, such as those present in the overlap zone, the MR-CB complex, or the lateral crest. Furthermore, it still remains a possibility that interactions between the disc and the actin cytoskeleton may also play a role in these movements. Overall, this work emphasizes the significance of disc contractility in establishing and maintaining parasite attachment, ultimately leading to key aspects of host pathology.

## Acknowledgements

This work was supported by a EuPATH/GiardiaDB award and a R01AI077571 award to SCD. We graciously thank the UC Davis MCB Microscopy Core for helpful advice on the SDC and SIM microscopes.

## Figure Legends

**Supplemental Figure S1. Movie of ventral disc contraction movements**

AVI file associated with image stack used in Figure 2B.

**Supplemental Figure S2. Movie of ventral disc compression movements**

AVI file associated with image stack used in Figure 2D.

**Supplemental Figure S3. Movie of ventral disc constriction movements**

AVI file associated with image stack used in Figure 2F.

**Supplemental Figure S4. Sequential knockout of all four DAP7268 alleles with antibiotic cassettes.**

Genomic DNA from partial or complete DAP7268 knockout strains and from WBC6 was PCR-amplified with primers 7268LeftF and 7268RightR, which bind outside the homology arms of the hygromycin (Hyg), neomycin (Neo) and blasticidin (Bsd) repair templates. Electroporation (EP) of the 7268Bsd repair template into the Cas9/7268gRNA908R expression strain resulted in partial knockout of DAP7268, with a 1.6 kb wild-type DAP7268 band and a 2.5 kb band indicating insertion of the 0.9 kb Bsd cassette into the gene. Quadruple knockout was achieved after a second electroporation of either the 7268Neo or the 7268Hyg repair templates into the 7268Bsd partial KO strain. The resulting quadruple KO strains lack the 1.6 kb wild-type DAP7268 band and have either a 2.9 kb or 3.1 kb band indicating insertion of the 1.3 kb Neo cassette or the 1.5 kb Hyg antibiotic cassette, respectively.

The same primers (7268LeftF and 7268RightR) were used to amplify the mutated DAP7268 locus for all alleles, and pooled PCR amplicons were sequenced with Nanopore long read sequencing (UC Berkeley). Reads corresponding to three clones of the DAP7268KO (A5, B2, C7) were mapped back to the WT DAP7268 locus indicate a gap in the area of the Cas9 DSB site and roughly 5-10 nucleotides flanking this region. No reads spanned the length of the WT locus using the BWA and Samtools software packages, and presentation using the Integrative Genome View (IGV) software shows the overall coverage of reads that and indicates complete insertion of cassettes into all four alleles. Reads from clones were also mapped back to the BSD and the HYG repair templates and coverage maps indicate the inserted region, with roughly 50% of alleles mapping back to either the BDS or HYG repair templates.

Lastly, mapping of reads to the 7268gRNA908R gRNA position indicates the 5 mutations made in the HDR templates to prevent Cas9-mediated DSB to mutated DAP7268 loci.

**Supplemental Figure S5. Screening of quad KO in 7268BH clones with expression strain and wild-type controls.**

Genomic DNA from 7268BH quadruple knockout clones (B2 through E11), the Cas9 and 7268gRNA908R expression strain (Cas) and WBC6 (Wt) was PCR-amplified with primers 7268LeáF and 7268RightR, which bind outside the homology arms of the hygromycin (Hyg) and blasàcidin (Bsd) repair templates. The 7268BH quadruple KO clones lack the 1.6 kb wild-type DAP7268 band and instead contain 3.1 kb and 2.5 kb bands indicaàng inseràon of the 1.5 kb hygromycin and 0.9 kb blasàcidin anàbioàc casseâes into all four copies of the DAP7268 gene.

**Supplemental Figure S6. Quadruple KO clones 7268BH B2 and quadKO plus knock-in of disc edge marker in clones 7268BH_12139mNG 3B11 and 3E9.**

Genomic DNA from DAP7268 quadruple knockout strains and WBC6 was PCR-amplified with primers 7268LeáF and 7268RightR, which bind outside the homology arms of repair templates used for knockout. All DAP7268 quadruple knockout strains lack the 1.6 kb wild-type DAP7268 band present in WBC6 and have a 2.5 kb band indicaàng inseràon of the 0.9 kb Bsd casseâe into the gene. Inseràon of either the hygromycin resistance casseâe (Hyg) or the hygromycin casseâe plus DAP12139-mNeonGreen, a marker of the disc edge, into the gene encoding DAP7268 results in 3.1 kb or 6.2 kb bands, respecàvely.

Similar nanopore sequencing of amplicons was performed for the DAP7268_DAP12139mNG strain for two clones (B11 and E9) as with the DAP7268KO strains. Again, coverage maps of reads mapping to the WT locus of each of these clones indicate a complete deletion of all WT alleles of the DAP7268locus; reads mapping to the BSD_DAP12139mNG and the HYG locus also indicate roughly 75% of reads mapping to the BSD cassette and 25% to the HYG_12139mNG cassette.

**Supplemental Figure S7. DAP7268 morpholino-based knockdowns have severe disc and lateral crest structural defects resulting in flattened, closed disc conformation.**

Negative staining of isolated discs from both wild type (A, D) and DAP7268KD (B,C,E) demonstrate disc body (body), and multiple overlap zone (oz) and multiple and extraneous regions of the lateral crest (lc) defects as compared to the wild type disc conformation. Other components of the microtubule cytoskeleton are noted, including the dense bands (db), the basal bodies (bb), the funis (fn) and the four flagellar pairs (AF = anterior, CF = caudal, VF = ventral, and PF = posteriolateral). Scale bars = 1 µm (A-C) or 200 nm (D).

**Supplemental Figure S8. Baumkuchen disc edge movie**

AVI file associated with image stack used in Figure 6A.

**Supplemental Figure S9. Wt compression of disc edge movie**

AVI file associated with image stack used in Figure 6C.

**Supplemental Figure S10. Wt constriction of disc edge movie**

AVI file associated with image stack used in Figure 6C.

**Supplemental Figure S11. 7268KO+ 12139 aberrant disc edge dynamics**

AVI file associated with image stack used in Figure 6C.

