## Supplementary material for "Dynamic ventral disc contraction is necessary for *Giardia* attachment and host pathology": S4

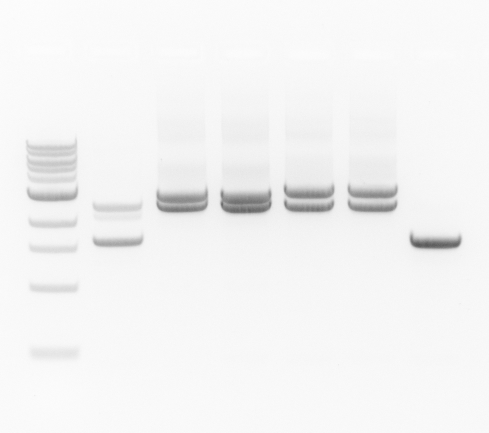

Bsd

Neo

EP1

Bsd

Neo

EP2

Bsd

Hyg

EP2

Bsd

Hyg

EP1

Bsd

Wt

Wt 1.6 kb

Bsd 2.5 kb

Hyg 3.1 kb

Neo 2.9 kb

Sequential knockout of DAP7268 with antibiotic cassettes. Genomic DNA from partial or complete DAP7268 knockout strains and from WBC6 was PCR-amplified with primers 7268LeftF and 7268RightR, which bind outside the homology arms of the hygromycin (Hyg), neomycin (Neo) and blasticidin (Bsd) repair templates. Electroporation (EP) of the 7268Bsd repair template into the Cas9/7268gRNA908R expression strain resulted in partial knockout of DAP7268, with a 1.6 kb wild-type DAP7268 band and a 2.5 kb band indicating insertion of the 0.9 kb Bsd cassette into the gene. Quadruple knockout was achieved after a second electroporation of either the 7268Neo or the 7268Hyg repair templates into the 7268Bsd partial KO strain. The resulting quadruple KO strains lack the 1.6 kb wild-type DAP7268 band and have either a 2.9 kb or 3.1 kb band indicating insertion of the 1.3 kb Neo cassette or the 1.5 kb Hyg antibiotic cassette, respectively.

Partial or complete KO of DAP7268 with Bsd plus Neo or Hyg
