## Supplementary material for "Dynamic ventral disc contraction is necessary for *Giardia* attachment and host pathology": S5

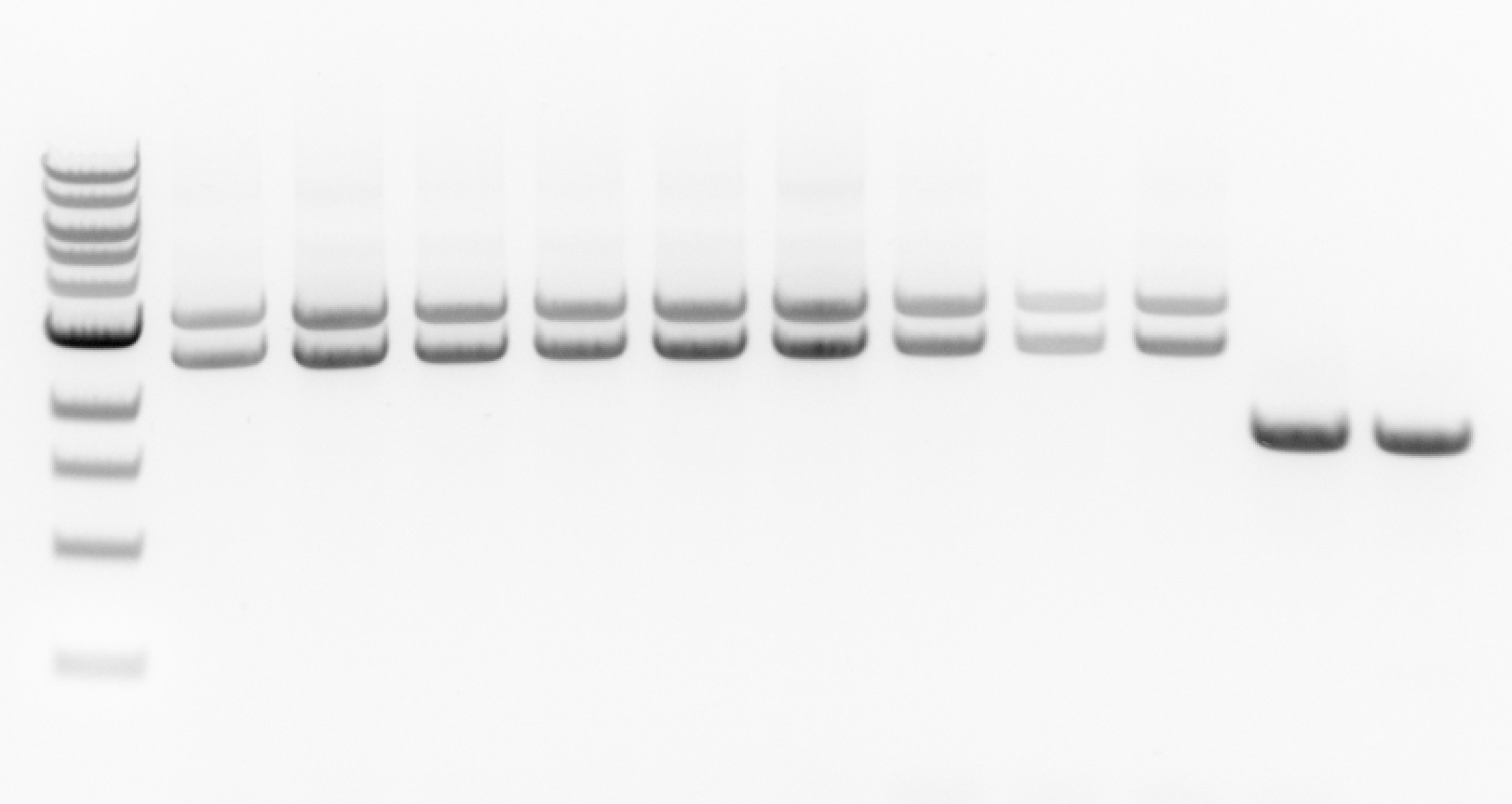

**B2**

C3

G4

**A5**

H6

**C7**

F8

B10

E11

Cas

Wt

Wt 1.6 kb

Bsd 2.5 kb

Hyg 3.1 kb

Genomic DNA from 7268BH quadruple knockout clones (B2 through E11), the Cas9 and 7268gRNA908R expression strain (Cas) and WBC6 (Wt) was PCR-amplified with primers 7268LeftF and 7268RightR, which bind outside the homology arms of the hygromycin (Hyg) and blasticidin (Bsd) repair templates. The 7268BH quadruple KO clones lack the 1.6 kb wild-type DAP7268 band and instead contain 3.1 kb and 2.5 kb bands indicating insertion of the 1.5 kb hygromycin and 0.9 kb blasticidin antibiotic cassettes into all four copies of the DAP7268 gene. Clones in bold (A5, B2, C7 were selected for long-read sequencing ).

7268BH clones with expression strain and wild-type controls

**DAP7268KO (clones A5, B2, C7) Nanopore long reads mapped to DAP7268 reference sequence**

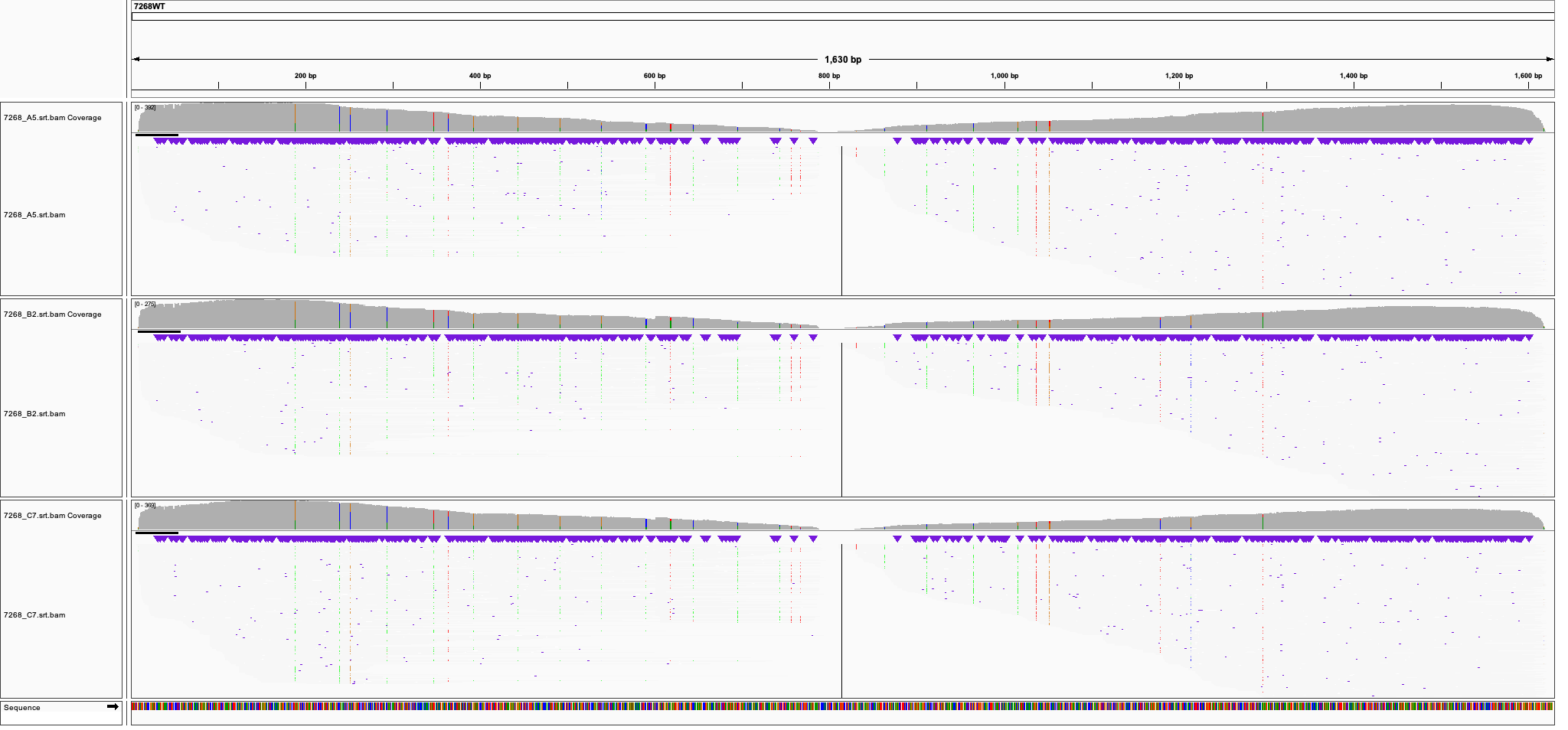

**DAP7268KO (clones A5, B2, C7) Nanopore long reads mapped to DAP7268_BSD cassette reference sequence**

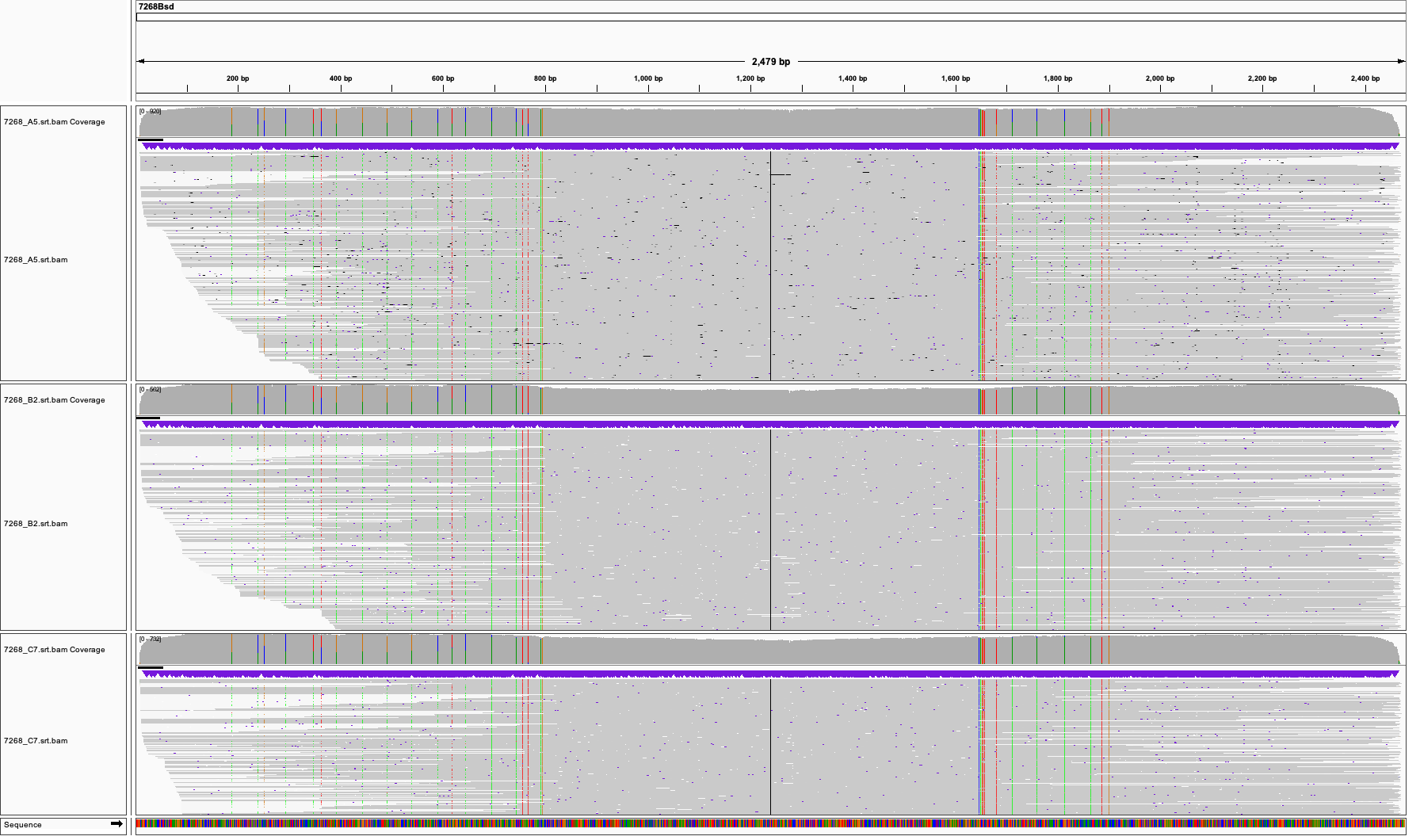

**DAP7268KO (clones A5, B2, C7) Nanopore long reads mapped to DAP7268_BSD cassette reference sequence**

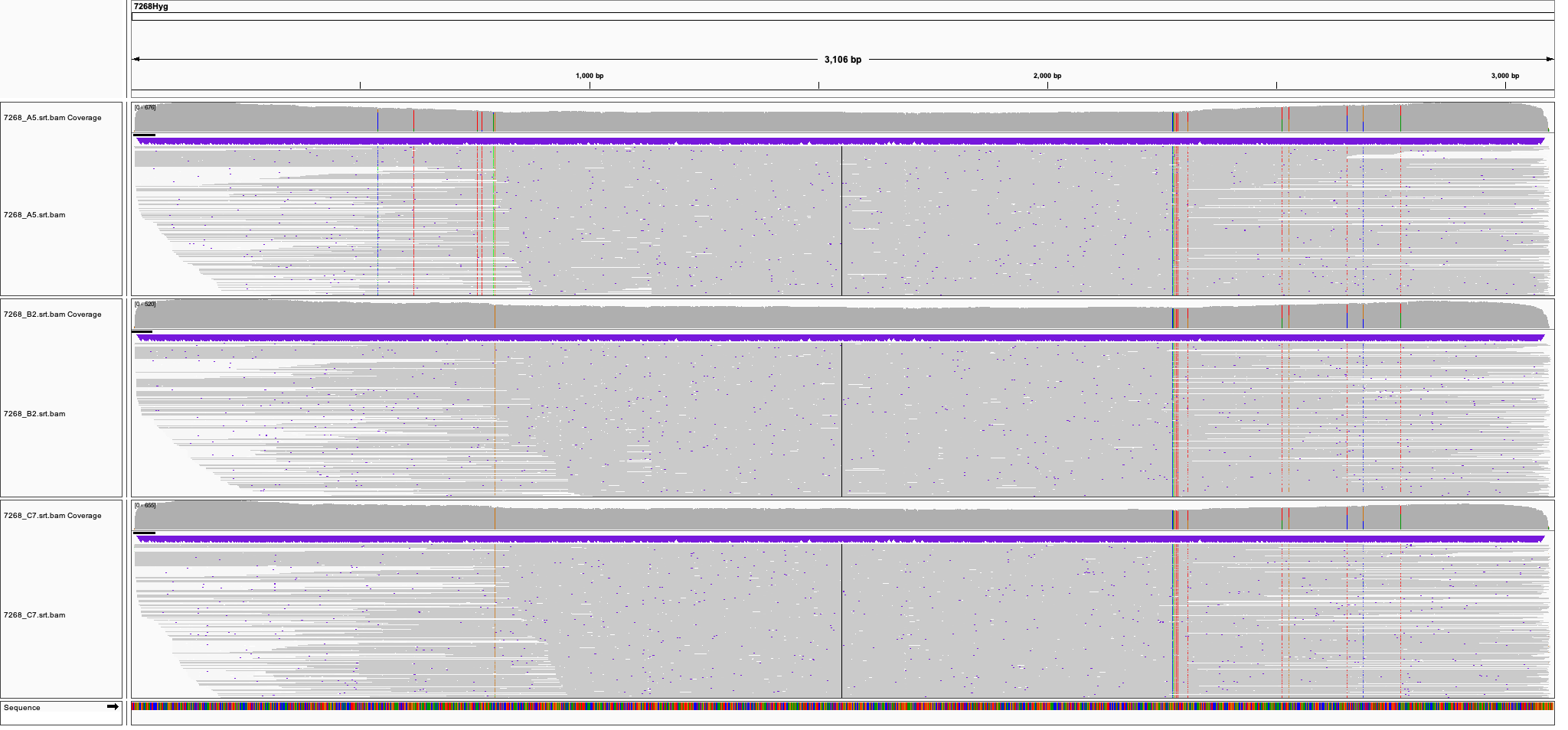

**DAP7268KO (clones A5, B2, C7) Nanopore long reads mapped to DAP7268_WT cassette indicated mutated PAM and gRNA region to prevent DSB in mutated DAP7268 locus**

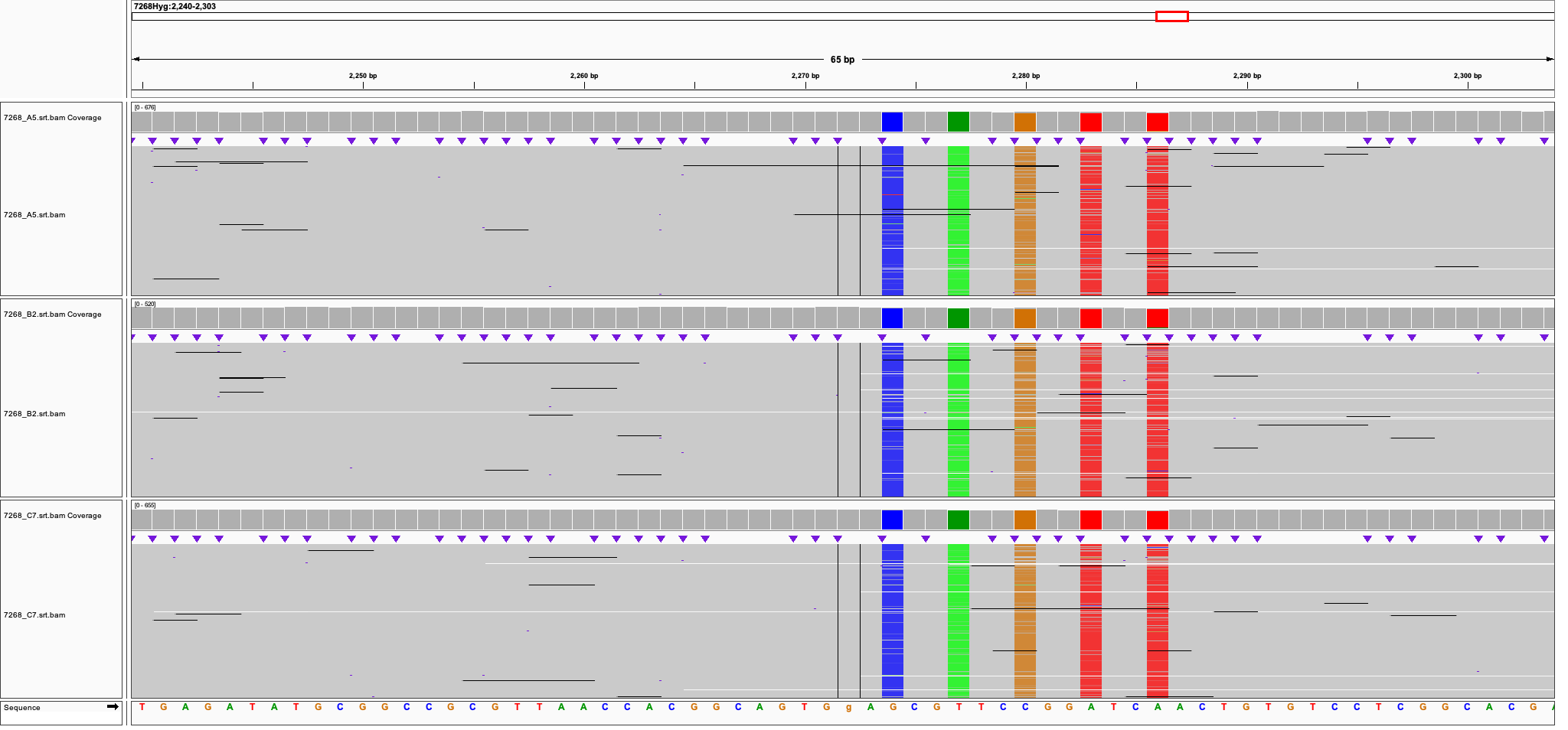

**______________________________** 7268gRNA908R
