## Supplementary material for "Dynamic ventral disc contraction is necessary for *Giardia* attachment and host pathology": S6

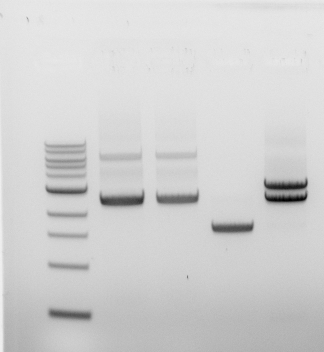

Wt 1.6 kb

Bsd 2.5 kb

Hyg 3.1 kb

12139mNGHyg 6.2 kb

Quadruple KO clones 7268BH B2 and 7268BH_12139mNG 3B11 and 3E9

3B11

3E9

WBC6

7268BH

7268BH_12139mNG

Genomic DNA from DAP7268 quadruple knockout strains and WBC6 was PCR-amplified with primers 7268LeftF and 7268RightR, which bind outside the homology arms of repair templates used for knockout. All DAP7268 quadruple knockout strains lack the 1.6 kb wild-type DAP7268 band present in WBC6 and have a 2.5 kb band indicating insertion of the 0.9 kb Bsd cassette into the gene. Insertion of either the hygromycin resistance cassette (Hyg) or the hygromycin cassette plus DAP12139-mNeonGreen, a marker of the disc edge, into the gene encoding DAP7268 results in 3.1 kb or 6.2 kb bands, respectively.

B2

**DAP7268KO_DAP12139 (clones B11, E9) Nanopore long reads mapped to DAP7268 reference sequence**

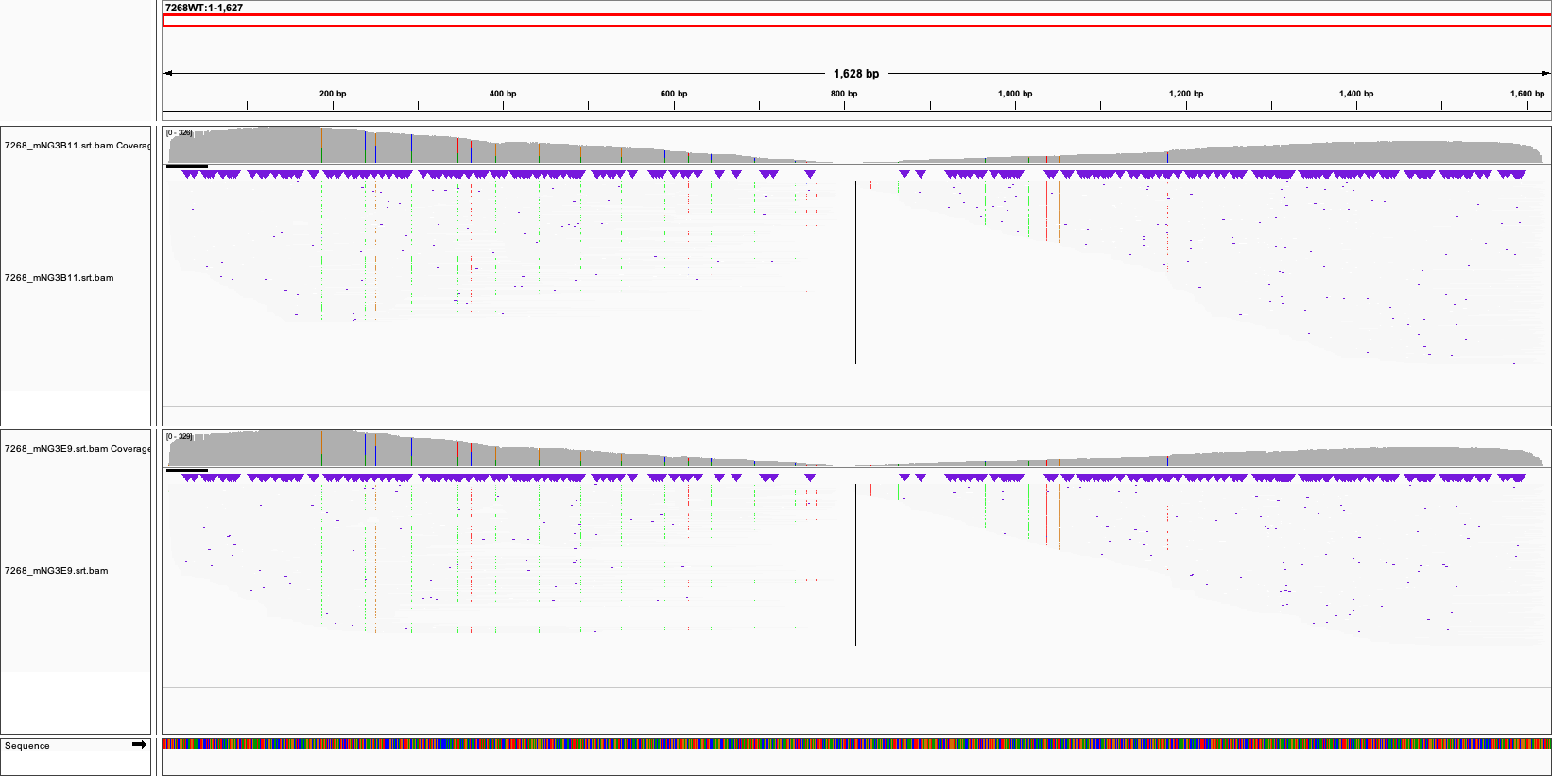

**DAP7268KO_DAP12139mNG (clones B11, E9) Nanopore long reads mapped to BSD_12139mNG reference sequence**

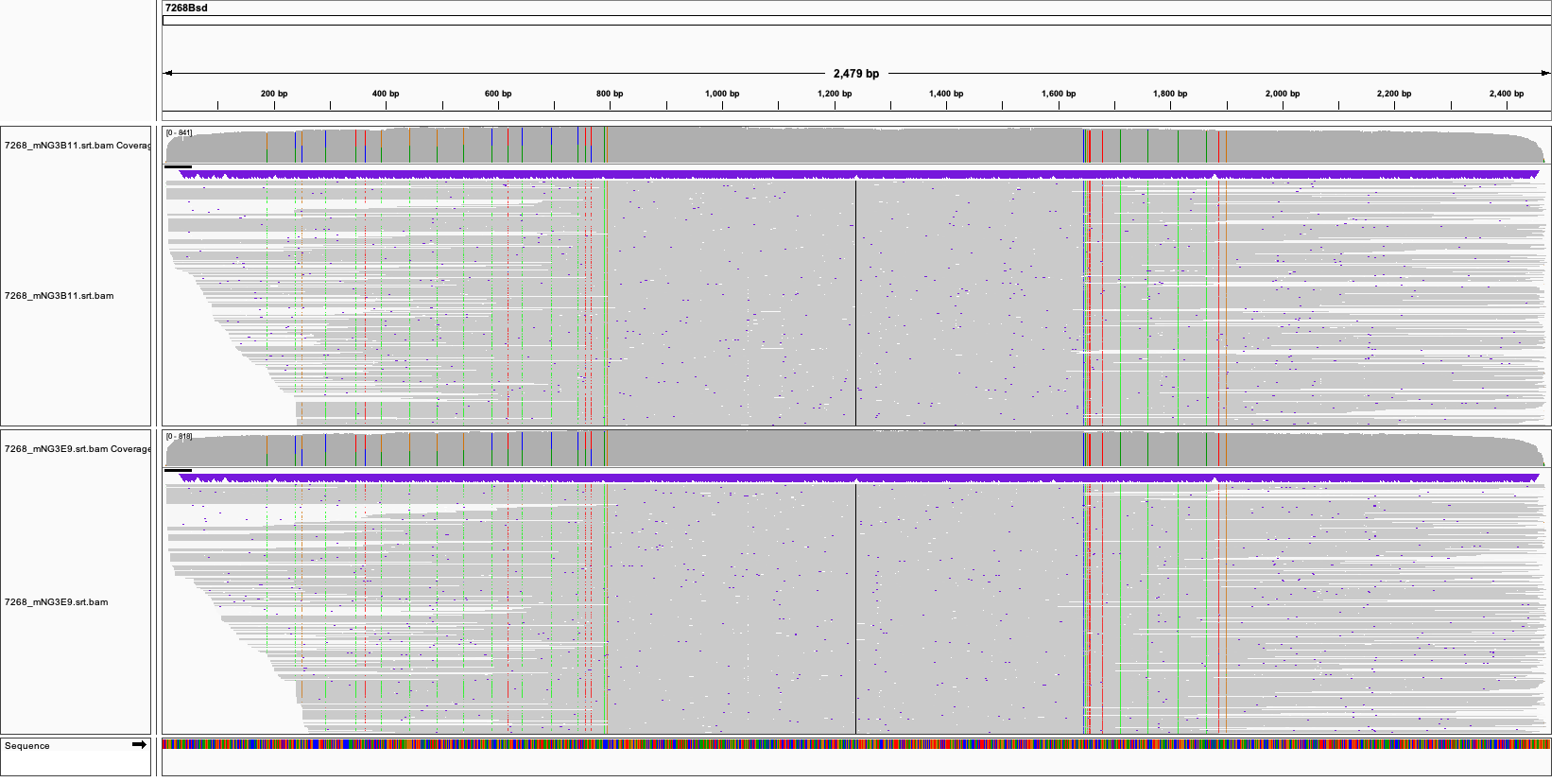

**DAP7268KO_DAP12139mNG (clones B11, E9) Nanopore long reads mapped to HYG reference sequence**

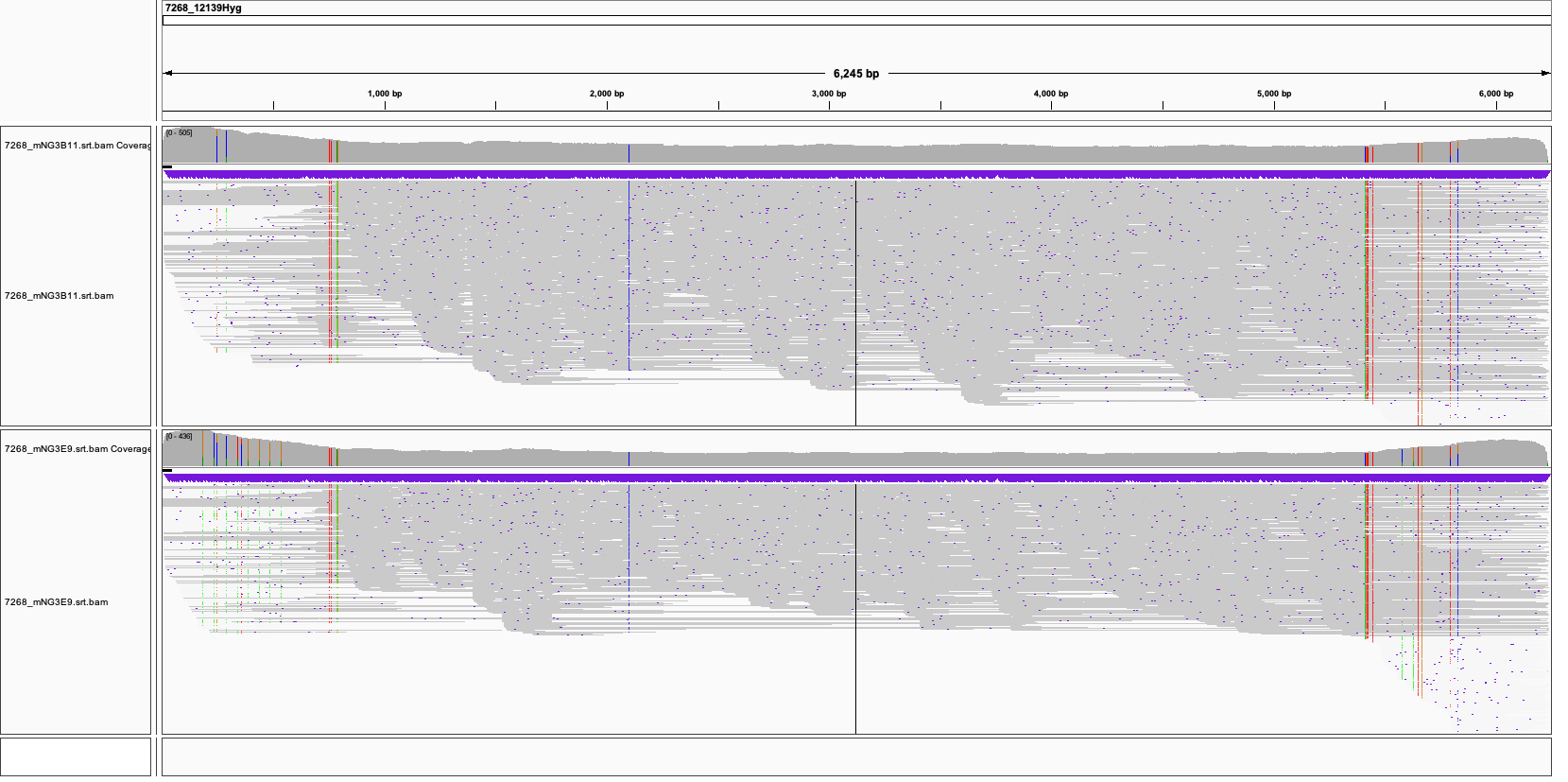
