## Supplementary figures and images for "Dynamic ventral disc contraction is necessary for *Giardia* attachment and host pathology"

### S7

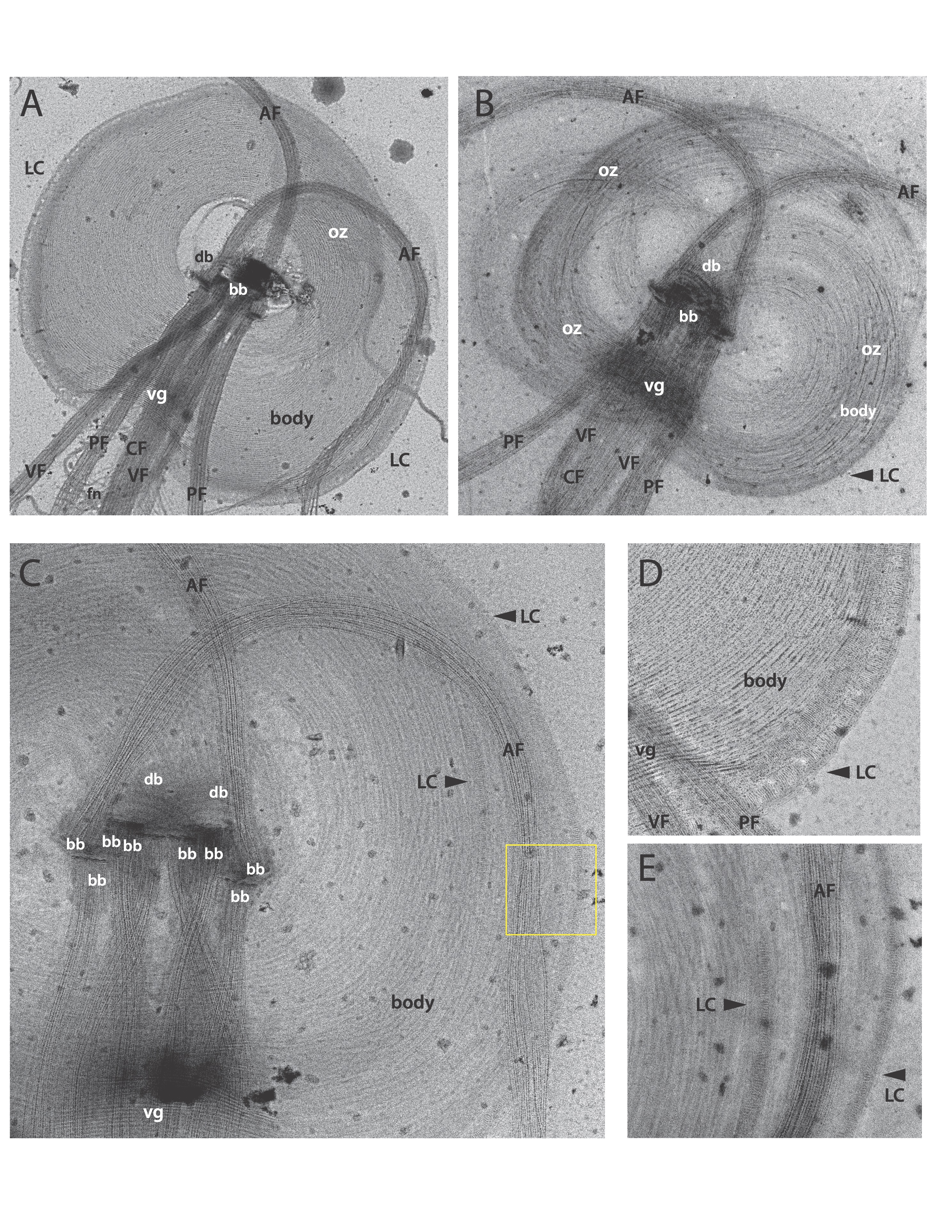
